# Allele-agnostic *KCNH1* suppression rescues gain-of-function developmental epileptic encephalopathy

**DOI:** 10.64898/2026.11.06.764929

**Authors:** Wanhao Chi, Erin E. Corbett, Christopher H. Thompson, Dina Simkin, Hiba Adil, Nicole A. Hawkins, Noah A. Callow, Levi Barse, Alfred L. George, Jennifer A. Kearney, Evangelos Kiskinis

## Abstract

Gain-of-function variants in *KCNH1*, which encodes the voltage-gated potassium channel Kv10.1^1,2^, cause developmental and epileptic encephalopathies (DEEs) characterized by seizures, developmental delay and intellectual disability^3–5^. While pathogenic variants enhance channel activity^6^, loss of *KCNH1* is tolerated in humans and mice^7,8^, suggesting that allele-agnostic gene suppression may represent a viable therapeutic strategy. Here, we developed a gapmer antisense oligonucleotide (ASO) targeting *KCNH1* and evaluated its efficacy in human induced pluripotent stem cell (iPSC)-derived neurons and a knock-in mouse model of *KCNH1*-related disorder. Neurons carrying the pathogenic *KCNH1* variants I494V or G375R exhibited a shared phenotype characterized by hyperpolarized resting membrane potential and abnormalities in neuronal network activity despite largely preserved intrinsic firing properties. ASOs efficiently reduced *KCNH1* expression without disrupting network activity in control neurons. In contrast, ASO treatment normalized network dysfunction across both mutant genotypes, demonstrating that distinct *KCNH1* gain-of-function variants converge on a reversible disease state. To assess the efficacy of ASOs *in vivo*, we generated a new *Kcnh1*-R357Q knock-in mouse model that exhibited profound seizures and early mortality. A single ASO treatment reduced seizure burden significantly and restored survival to near-normal levels. Together, these findings establish *KCNH1* suppression as a precision therapeutic strategy for *KCNH1*-related DEEs and provide a framework for treating gain-of-function epilepsies in which partial gene reduction is clinically tolerated.

## Main

Developmental and epileptic encephalopathies (DEEs) are among the most severe neurological disorders of childhood, characterized by early-onset, drug-resistant seizures, developmental impairment, and intellectual disability^9^. Most DEEs have a genetic basis, with pathogenic variants identified in hundreds of genes, particularly those encoding ion channels^10–12^. Although both gain-of-function (GoF) and loss-of-function (LoF) variants can cause disease in many ion channel genes^13^, a subset of DEEs is driven predominantly by GoF variants, whereas partial loss of channel function is relatively well tolerated^7^. This genetic architecture creates an opportunity for allele-agnostic gene suppression using gapmer antisense oligonucleotides (ASOs), enabling a single therapeutic strategy to target multiple pathogenic variants that share a common GoF mechanism. Here, we provide proof of concept that allele-agnostic gapmer ASO-mediated suppression of *KCNH1* can therapeutically target distinct GoF variants causing *KCNH1*-related DEEs. Partial suppression of *KCNH1* was well tolerated in control human neurons, rescued neuronal network dysfunction caused by distinct GoF variants, and markedly reduced seizures and mortality in a novel knock-in mouse model. These findings establish *KCNH1* suppression as a potential therapeutic strategy for *KCNH1*-related DEEs and provide proof of concept for allele-agnostic treatment of GoF neurodevelopmental disorders in which partial gene reduction is tolerated.

*KCNH1* encodes K_V_10.1 (Eag1), the founding member of the EAG family of voltage-gated potassium channels^1,2^. Genetic variants in *KCNH1* cause rare DEEs including Temple-Baraitser Syndrome (TBS) and Zimmermann-Laband Syndrome (ZLS), characterized by infantile- or childhood-onset seizures, developmental delay, and intellectual disability^3–5,14^. To date, approximately 50 affected individuals carrying more than 20 unique pathogenic variants have been reported (Fig. 1a and Extended Data Fig. 1). Most pathogenic variants arise *de novo* and cluster within conserved functional domains of Kv10.1. Electrophysiological studies in heterologous expression systems demonstrate that disease-associated variants result in GoF effects, enhancing channel function through altered voltage dependence and gating kinetics, and leading to hyperpolarized resting membrane potentials (Fig. 1b and Extended Data Fig. 2)^6^. Importantly, genetic analysis shows that *KCNH1* is among the most missense-constrained genes in the general population, with a missense constraint percentile above the 99^th^ percentile, indicating strong selection against amino acid-altering variation (gnomAD; missense Z-score=5.57)^7^ (Fig. 1c, top). In contrast, *KCNH1* exhibits a substantially lower constraint against predicted LoF variants (LOEUF = 0.54)^7^ (Fig. 1c, bottom). These population genetic data are consistent with pathogenic *KCNH1* variants acting predominantly through altered or enhanced channel function rather than haploinsufficiency. The plausibility that GoF mechanisms drive disease suggest that inhibiting *KCNH1* channel function may be therapeutically beneficial. However, direct pharmacologic inhibition of Kv10.1 is challenging due to high homology with the cardiac potassium channel KCNH2 (Kv11.1, hERG)^2^, creating substantial risk for cardiac toxicity with conventional small-molecule approaches^5,15^. We therefore developed a gapmer-based ASO therapeutic strategy designed to reduce *KCNH1* expression.

**Fig. 1:**
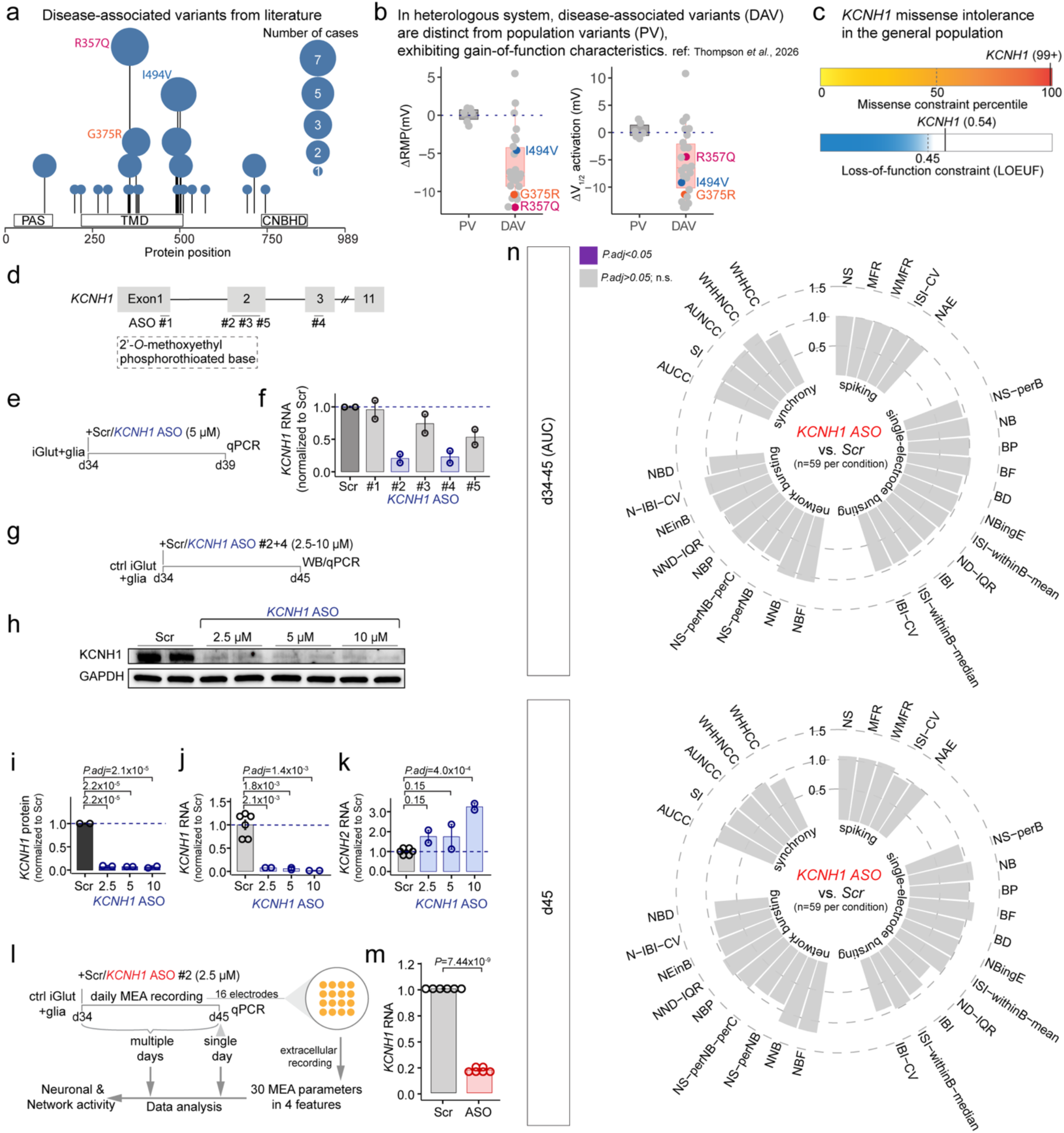
Human genetics of *KCNH1* and development of *KCNH1*-targeting gapmer ASOs. **a,** Distribution of pathogenic *KCNH1* variants reported in the literature across the KCNH1 protein. Variants examined in this study are labelled. Dot size indicates the number of affected individuals carrying each variant. PAS, Per-Arnt-Sim domain; TMD, transmembrane domain comprising the S1-S4 voltage-sensing domains and the S5-pore-S6 pore domain; CNBHD, cyclic nucleotide-binding homology domain. **b,** Functional effects of disease-associated variants (DAVs) and population variants (PVs) reported in a previous heterologous expression study^6^. Resting membrane potential (RMP), and half-activation voltage (V_1/2_) are shown relative to wild-type (WT) channels. **c,** Population constraint metrics for missense and loss-of-function variation in *KCNH1* based on Genome Aggregation Database (gnomAD) data^7^. LOEUF: loss-of-function observed/expected upper bound fraction. **d,** Location of ASOs within the schematic *KCNH1* genomic sequence. ASOs were incorporated 2’-*O*-methoxyethyl (2′-MOE) sugar modifications and phosphorothioate backbone**. e,** Experimental timeline for ASO screening. **f,** *KCNH1* expression following treatment with scrambled control (Scr) or various *KCNH1*-targeting ASOs. Data were normalized to *GAPDH* and *GPI* for each sample and then to Scr. **g,** Experimental timeline for ASO dose-response experiments. ASO-2 and ASO-4 were mixed at 1:1 ratio and the final dose for each is half of the total. **h,** Western blot analysis following treatment with Scr (10 μM) or various concentrations of *KCNH1*-targeting ASO. **i,** Quantification of KCNH1 protein from panel h. Data were normalized to GAPDH for each sample and then to Scr. n=2 per condition. Statistical comparisons were performed using one-way ANOVA followed by Tukey’s post hoc test. **j,k,** *KCNH1* and *KCNH2* RNA levels following treatment with Scr or various concentrations of *KCNH1*-targeting ASO. Data were normalized to *GAPDH* and *GPI* for each sample and then to Scr. n=2-6 per condition. Statistical comparisons were performed using one-way ANOVA followed by Tukey’s post hoc test. **l,** Schematic of experimental timeline for ASO treatment and MEA recordings, and analysis workflow. **m,** *KCNH1* expression following treatment with Scr or *KCNH1*-targeting ASO. n=6 per condition from six independent differentiation batches. Statistical comparisons were performed using Welch’s two-sample *t*-test. **n,** Partial *KCNH1* knockdown does not alter spontaneous neuronal network activity in control neurons measured by MEA. Normalized data to Scr were grouped into four functional categories. For days 34-45, the area under the curve (AUC) was calculated for each well; raw values from day 45 were analyzed. n=59 wells per condition from six independent differentiation batches. Statistical comparisons were performed using Wilcoxon rank-sum test, followed by Benjamini-Hochberg correction for multiple comparisons across the 30 parameters. See Supplemental Table 2 for the full name of each abbreviation.

Given that *KCNH1* LoF is relatively tolerated in humans and that the *Kcnh1* knockout mice are viable without gross developmental abnormalities^8^, we designed allele-agnostic -rather than allele-specific-ASOs targeting the first 3 exons of the mRNA (Fig. 1d and Supplemental Table 1). The ASOs were 20 nucleotides in length and incorporated 2’-*O*-methoxyethyl (2′-MOE) sugar modifications and phosphorothioate backbone, chemical modifications that are also used in FDA-approved ASOs^16^. To assess target engagement, we screened ASOs in human control induced pluripotent stem cell (iPSC)-derived excitatory (iGlut) neurons at day 34 of differentiation (Fig. 1e and Extended Data Fig. 3a). At this stage, *KCNH1* expression had plateaued and neurons had established functional networks, as indicated by robust expression of the pre- and postsynaptic markers SYN1 and PSD95, and detection of spontaneous activity across all 16 electrodes in multi-electrode array (MEA) recordings (Extended Data Fig. 3b-d). Initial screening identified three ASOs that reduced *KCNH1* transcript levels, including two lead candidates, ASO-2 and ASO-4, that reduced *KCNH1* mRNA by approximately 70% relative to scrambled controls after 5 days (Fig. 1f). Dose-response analyses of combined ASO-2 and ASO-4 treatment (2.5-10 µM) showed a dramatic reduction in *KCNH1* mRNA and protein levels (Fig. 1g-j and Extended Data Fig. 4). These ASOs did not reduce expression of the closely related paralog *KCNH2*, supporting selective suppression of *KCNH1* (Fig. 1k). Lastly, treatment of control iGlut neurons with ASO-2 (2.5 µM) substantially reduced *KCNH1* expression (Fig. 1l,1m), without significantly altering any measure of spontaneous activity assessed by MEA recordings (P.adj >0.05; Fig. 1n), indicating that *KCNH1* knockdown is tolerated at the neuronal network level.

To model *KCNH1*-DEE in human neurons, we used precise CRIPSR/Cas9 genome editing to introduce two recurrent, GoF disease-associated *KCNH1* variants (I494V and G375R; Fig. 1a) into a control iPSC line (Fig. 2a). The resulting isogenic pairs of iPSC lines exhibited normal karyotypes, expressed markers of pluripotency, maintained a normal *KCNH1* copy number, and harbored no mutations at predicted off-target sites (Extended Data Figs. 5-6). Importantly, neither disease-associated variant affected the efficiency of differentiation into iGlut neurons (Fig. 2b) or altered KCNH1 protein abundance (Fig. 2c and Extended Data Fig. 7). To assess whether the variants affected channel trafficking, we generated mCherry-tagged *KCNH1* expression constructs. Whole-cell voltage-clamp recordings in HEK293T cells showed that the tagged channels largely retained the properties of untagged channels (Extended Data Fig. 8a-b). Cell surface biotinylation showed that both mutant proteins localized to the plasma membrane at levels comparable to wild-type KCNH1 (Fig. 2d and Extended Data Fig. 8c).

**Fig. 2:**
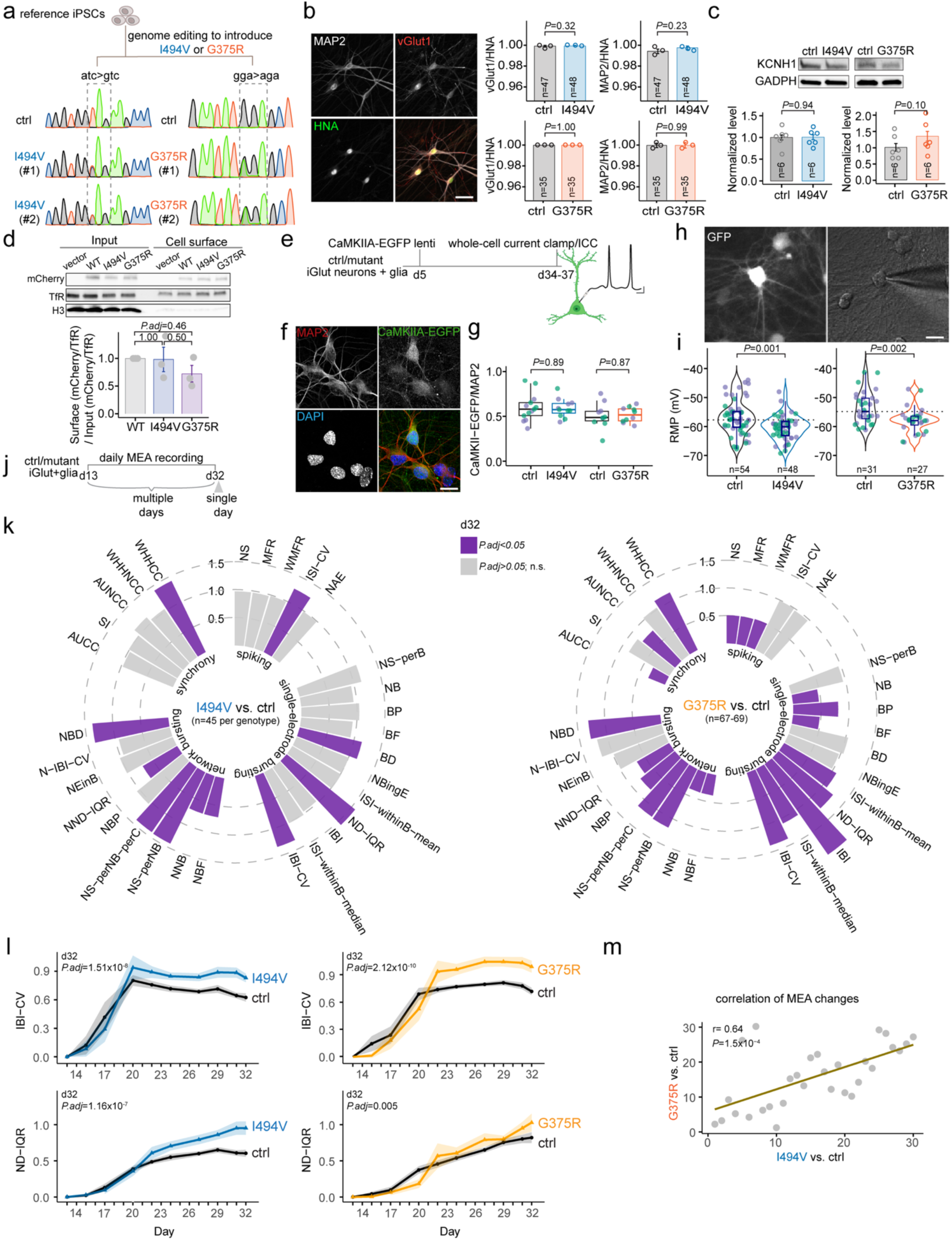
*KCNH1* gain-of-function variants hyperpolarize the resting membrane potential and disrupt neuronal network activity in human iPSC-derived neurons. **a,** Generation and validation of isogenic KOLF2.1J iPSC lines carrying the pathogenic *KCNH1* variants I494V or G375R. Variants are named according to transcript isoform 1 (NCBI RefSeq NM_172362). Two independent clones were generated for each variant. Clone 1 from each mutant was used in the study. **b,** Comparable differentiation efficiency of mutant and control iPSCs into glutamatergic (iGlut) neurons. vGlut1, MAP2, and human nuclear antigen (HNA) were used as markers of glutamatergic neurons, neurons, and human cells, respectively. Scale bar, 25 μm. n=35-48 images per condition from three independent differentiation batches. Statistical comparisons were performed using Welch’s two-sample *t*-test**. c,** Comparable KCNH1 protein expression in mutant and control iGlut neurons. n=6 samples per condition from two independent differentiation batches. Statistical comparisons were performed using Welch’s two-sample *t*-test**. d,** Comparable cell-surface expression of mCherry fused I494V and G375R mutant Kv10.1 channels relative to WT following heterologous expression in HEK293T cells. Statistical comparisons were performed using one-way ANOVA followed by Tukey’s post hoc test. n=3 independent transfection. Transferrin receptor (TfR) and histone H3 (H3) were used as a positive and negative cell-surface protein marker, respectively. **e,** Experimental timeline for whole-cell current-clamp recordings. A CaMKIIA-EGFP lentiviral reporter was used to identify mature neurons. **f**, Representative immunofluorescence images showing expression of CaMKIIA-EGFP in iGlut neurons. MAP2 and DAPI were used as markers of glutamatergic neurons and nuclei. Scale bar, 20 μm. **g,** Comparable proportions of EGFP-positive neurons in control and mutant cultures. n=6 images per condition from two independent differentiation batches. Statistical comparisons were performed using Welch’s two-sample *t*-test**. h,** Representative fluorescence and DIC images of patched neurons. Scale bar, 20 μm. **i,** Neurons carrying the I494V or G375R variant exhibited significantly hyperpolarized resting membrane potentials (RMPs) relative to control neurons. n=27-54 neurons per condition from two independent differentiation batches. Statistical comparisons were performed using Welch’s two-sample *t*-test**. j**, Experimental timeline for MEA recordings. **k**, Normalized AUC data were grouped into four functional categories. n=45-69 wells per condition from three to four independent differentiation batches. Statistical comparisons were performed using Wilcoxon rank-sum test, followed by Benjamini-Hochberg correction for multiple comparisons across 30 parameters. **l**, Time course of normalized interburst interval coefficient of variation (IBI-CV) and normalized interquartile range duration (ND-IQR) in control and mutant neurons. Ribbons indicate 95% confidence intervals. Only non-feeding days were plotted. **m**, Correlation analysis of ranked MEA parameters between I494V and G375R neurons. Statistical significance was assessed using Spearman’s rank correlation test.

To investigate the functional consequences of disease-associated *KCNH1* variants, we performed whole-cell current-clamp recordings in iGlut neurons at days 34-37 of differentiation (Fig. 2e). To reduce variability associated with neuronal maturation, we labeled mature excitatory neurons using a CaMKIIα-EGFP reporter (Fig. 2f-g) and examined the intrinsic electrophysiological properties of EGFP-positive neurons. Consistent with previous studies in heterologous expression systems (Fig. 1b)^6^, and the established function of KCNH1 channels, neurons carrying the I494V or G375R variant exhibited significantly hyperpolarized resting membrane potentials (RMPs) compared to their respective isogenic controls (I494V: −60.45 ± 0.61 versus −57.02 ± 0.80 mV, *P* = 0.001; G375R: −57.92 ± 0.83 versus −53.68 ± 0.97 mV, *P* = 0.002; mean ± s.e.m.) (Fig. 2h-i). In contrast, neither variant significantly altered other intrinsic action potential (AP) properties including threshold, amplitude, half-width, depolarization and repolarization velocity, or input resistance (Extended Data Fig. 9). We also did not detect any differences in the fast component of the afterhyperpolarization (AHP), or evoked AP firing (Extended Data Figs. 9-10).

We next examined spontaneous neuronal network activity longitudinally using MEA recordings of iGlut neurons from day 13 to 32 (Fig. 2j). At day 32, both I494V and G375R neurons showed abnormalities in single-electrode and network bursting (Fig. 2k). Specifically, mutant neurons exhibited increased variability in the timing and duration of single-electrode bursts, reflected by an increased coefficient of variation of the interburst interval (IBI-CV) and an increased interquartile range of normalized burst duration (ND-IQR), respectively, suggesting temporal disorganization of bursting. These single-electrode bursting phenotypes were also evident longitudinally from days 13-32, indicating that they emerged during neuronal maturation and persisted thereafter (Fig. 2l, Extended Data Fig. 11a). Notably, the effects of I494V and G375R across MEA parameters were significantly concordant in both magnitude and direction at day 32 (r = 0.64) (Fig. 2m), as well as when considering the entire day 13-32 trajectory (r = 0.9) (Extended Data Fig. 11b), demonstrating that two distinct *KCNH1* GoF variants converge on a shared pattern of neuronal network dysfunction.

Having established that disease-associated *KCNH1* variants disrupt neuronal network activity, we next tested if reducing *KCNH1* expression could reverse these phenotypes. I494V and G375R neurons were treated at day 34 with a single dose of *KCNH1*-targeting ASO-2 (2.5 µM) or a scrambled control ASO, followed by MEA recordings (Fig. 3a). Neurons derived from both isogenic control lines were treated with the scrambled ASO and included as references. At day 45, ASO-2 reduced *KCNH1* expression by approximately 70% in both mutant lines (Fig. 3b), and MEA recordings showed that scrambled ASO-treated I494V and G375R neurons exhibited shared abnormalities in the regularity of bursting, including increased IBI-CV and ND-IQR relative to their respective isogenic controls (Extended Data Fig. 12a). Longitudinal analysis revealed that ASO-2 progressively normalized both measures, with values at day 45 no longer significantly different from isogenic controls (Fig. 3c-f and Extended Data Fig. 12b). Consistent with this rescue, comparison across the broader multidimensional MEA profile revealed no significant differences between ASO-2-treated mutant neurons and their respective scrambled ASO-treated isogenic controls (Fig. 3g-h). Together, these findings demonstrate that partial *KCNH1* suppression reverses shared network abnormalities caused by two distinct disease-associated GoF variants.

**Fig. 3:**
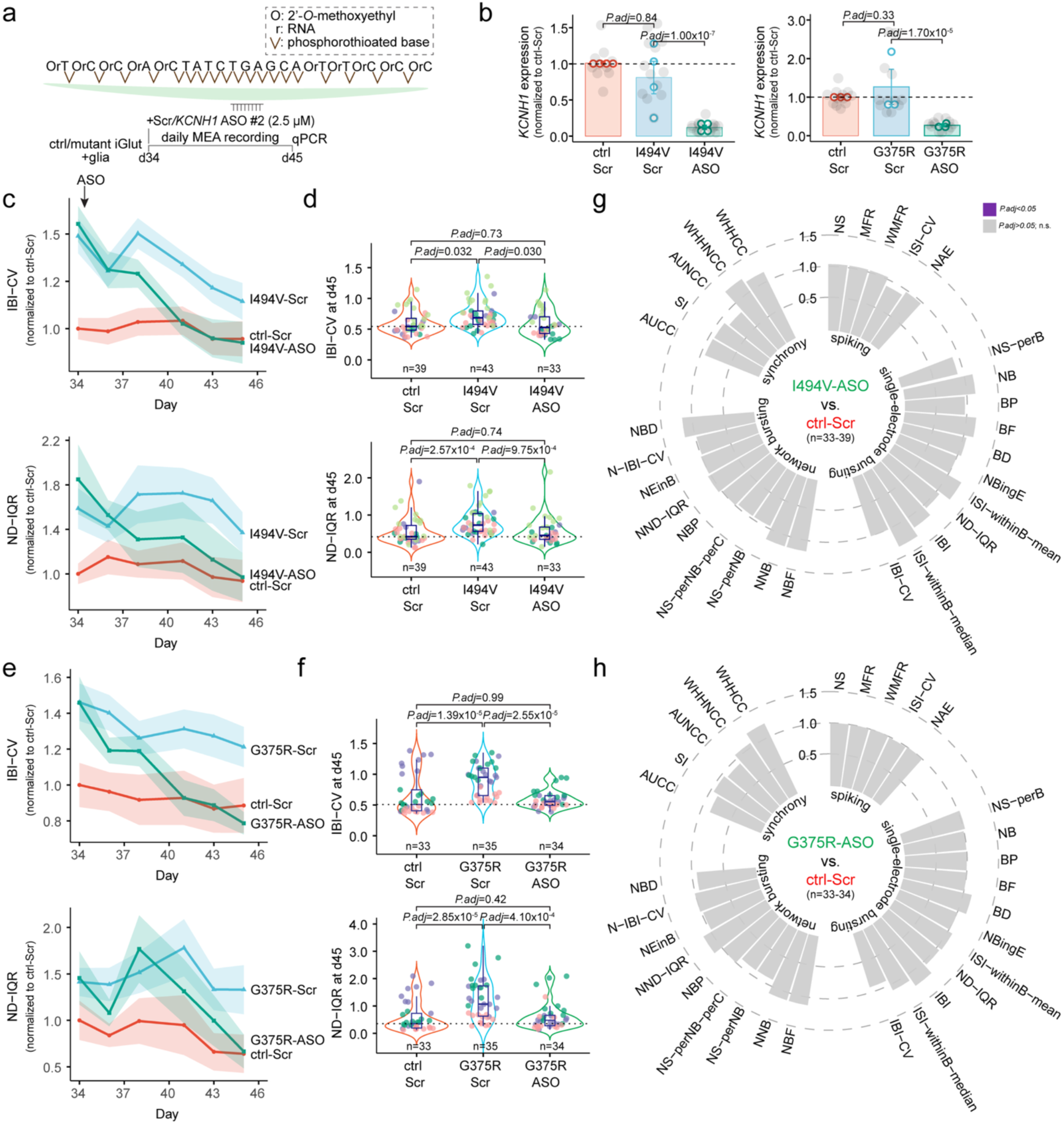
*KCNH1*-targeting ASOs normalize network abnormalities in *KCNH1* mutant neurons. **a,** Experimental timeline, ASO sequences, and chemical modifications. **b,** *KCNH1* expression in control or mutant neurons following treatment with Scr or *KCNH1*-targeting ASOs. n=10-13 samples per I494V condition from four intendent differentiation batches and n=10-12 samples per G375R condition from three intendent differentiation batches. Statistical comparisons were performed using one-way ANOVA followed by Tukey’s post hoc test. **c,e,** Time course of normalized interburst interval coefficient of variation (IBI-CV) and normalized interquartile range duration (ND-IQR) in control and mutant neurons treated with Scr or *KCNH1*-targeting ASOs. Ribbons indicate 95% confidence intervals. **d,f,** Raw d45 values of IBI-CV and ND-IQR. **g,h,** ASO treatment normalized MEA phenotypes in I494V (g) and G375R (h) mutant neurons to control levels. Normalized d45 data are shown. n=33-39 wells per I494V condition from four intendent differentiation batches (g) and n=33-34 wells per G375R condition from three intendent differentiation batches (h). For each MEA parameter, genotype or treatment effects were assessed using Kruskal-Wallis test, followed by Benjamini-Hochberg correction across 30 parameters. Parameters with significant effects were further analyzed using Dunn’s post hoc test for pairwise comparisons. See Extended Data Fig. 12 for the plots related to genotype difference and treatment effect.

We next asked whether *KCNH1* suppression could ameliorate disease phenotypes *in vivo* using a *Kcnh1* knock-in mouse model. Although human and mouse *KCNH1* protein sequences are highly conserved (97% amino acid identity; Extended Data Fig. 13a), the ASO-2 target sequence differs by two nucleotides between species (Extended Data Fig. 13b). We therefore generated a mouse-specific version of ASO-2 (mASO-2) (Extended Data Fig. 13c) and first evaluated target engagement in primary mouse cortical neurons. A single treatment with mASO-2 (2.5 or 5 µM) reduced *Kcnh1* transcript levels by more than 80% after 7 days compared to a scrambled ASO, demonstrating efficient suppression of endogenous mouse transcript (Extended Data Fig. 13d-e). To generate a mouse model of *KCNH1*-associated DEE we introduced the recurrent pathogenic, GoF R357Q variant into the endogenous mouse *Kcnh1* locus using CRISPR/Cas9 genome editing (Figs. 1a, 4a, and Extended Data Fig. 14a). PCR and Sanger sequencing confirmed presence of the desired mutation and absence of mutations at predicted off-target sites (Fig. 1a and Extended Data Fig. 14a-d). The R357Q variant did not alter *Kcnh1* mRNA or protein levels (Extended Data Fig. 14e-g). Heterozygous *Kcnh1^R357Q/+^*mice, hereafter referred to as *Kcnh1*^Q/+^, were born at expected Mendelian ratios but had significant mortality in the fourth week of life (Fig. 4b), coincident with convulsive behavioral seizures that were confirmed by electroencephalographic (EEG) recording (Fig. 4c). To maintain the mouse line, *Kcnh1^Q/+^*mice were treated with the investigational sodium channel inhibitor GS967 that improves survival but does not provide seizure freedom (Extended Data Video 1). Intracerebroventricular (ICV) administration of mASO-2 (10-30 µg) to wild-type (WT) pups at postnatal day 1 (P1) resulted in significant and durable reduction in *Kcnh1* levels by more than 30% at 4-12 weeks of age (Figs. 4d-f). To examine whether *Kcnh1* suppression could mitigate seizures and prevent lethality of *Kcnh1*^Q/+^ mice, we administered mASO-2 by ICV injection in P1 pups (Fig. 4g). To specifically determine the impact of mASO-2, we removed GS967 treatment after P24. A single neonatal dose of mASO-2 (30 µg) dramatically improved survival and markedly lowered seizure burden (Fig. 4h-i). *Kcnh1*^Q/+^ mice treated with mASO-2 had 12-week survival of 78% compared to <10% for vehicle-treated mice. Seizures were quantified in a 6-hour window overlapping the period of highest mortality for vehicle-treated or untreated *Kcnh1*^Q/+^ mice. Over the six-hour observation window, one mASO-2-treated mouse (1/12) exhibited a single convulsive seizure (Extended Data Video 2), while 90% (18/20) of vehicle-treated mice exhibited at least one seizure and 75% (15/20) had multiple seizures (Fig. 4i and Extended Data Video 3). Seizure frequency for vehicle-treated mice was similar to *Kcnh1*^Q/+^ mice that received no ICV treatment and were withdrawn from GS967 at weaning (Fig. 4i and Extended Data Video 4). Together, these findings demonstrate that a single neonatal dose of *Kcnh1*-targeting mASO-2 markedly reduces seizures and confers a sustained survival benefit in a mouse model of *Kcnh1*-associated DEE, supporting *KCNH1* suppression as a promising therapeutic strategy.

**Fig. 4:**
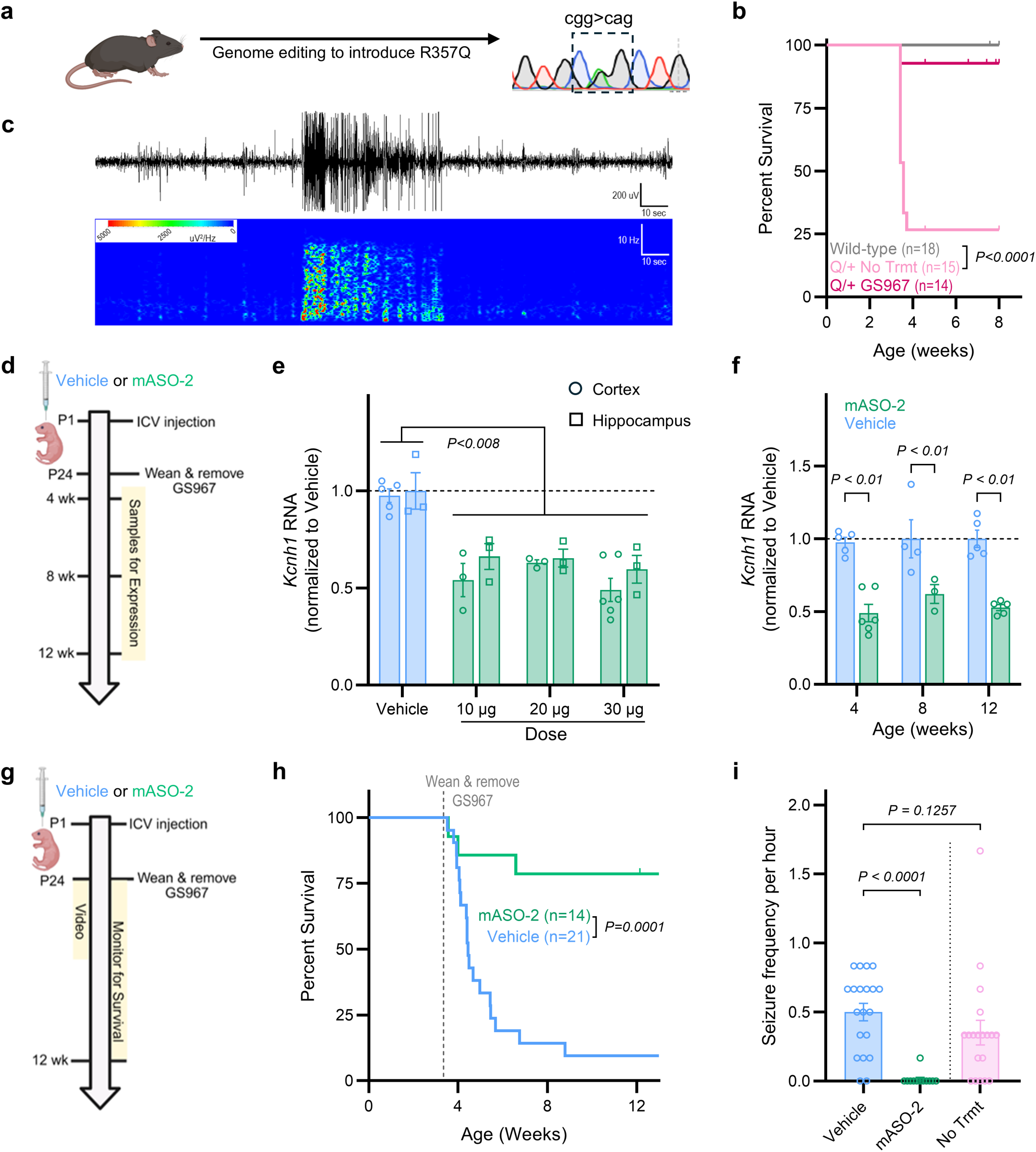
In vivo administration of mASO-2 reduces *Kcnh1* expression, improves survival and lowers seizure burden in a novel *Kcnh1^R357Q^* mouse model. **a,** Generation of a pathogenic *Kcnh1* variant corresponding to the human R357Q mutation on the C57BL/6J background. **b,c,** Heterozygous *Kcnh1^Q/+^*(Q/+) mice exhibit electrographic seizures with behavioral correlates (b), and early lethality that can be attenuated by treatment with GS967 (c). All *Kcnh1^Q/+^* mice received GS967 before weaning and were subsequently maintained on either GS967-containing chow (GS967) or standard chow (No Trmt). **d,** Experimental timeline for expression studies following a single intracerebroventricular (ICV) injection of mASO-2 at postnatal day 1 (P1). **e,** mASO-2 doses of 10-30 µg reduced *Kcnh1* transcript expression in hippocampus (squares) and cortex (circles) at P24 as quantified by RT-ddPCR. **f,** mASO-2 administration (30 µg) provided durable attenuation of *Kcnh1* expression for up to 12 weeks. **g,** Experimental timeline for efficacy studies. **h,** Kaplan-Meier survival curves of *Kcnh1^Q/+^* mice following a single neonatal ICV administration of mASO-2 (30 µg) or vehicle (n=14 mASO-2; n=21 vehicle). Treatments were compared using Mantel-Cox Log Rank test. **i,** Seizure frequency quantified in a 6-hour window overlapping the peak mortality window (n=12 mASO-2; n=20 vehicle). Seizure frequency quantified in the 6-hour peak mortality window for the ‘No Trmt’ group in panel b is provided for comparison (n=19). Seizure frequencies were compared using Kruskal-Wallis test, with Dunn’s post-hoc comparisons.

## Discussion

ASO-based therapies are promising approaches for DEEs, with preclinical studies demonstrating therapeutic efficacy across both GoF and LoF disease mechanisms^17–22^. Here, we demonstrate that non-allele specific suppression of *KCNH1* rescues neuronal network dysfunction caused by distinct pathogenic GoF variants and markedly improves seizures and survival *in vivo*. Importantly, these findings establish a therapeutic strategy that does not require development of variant-specific ASOs. When partial reduction of gene function is tolerated, a single allele-agnostic ASO may be applicable across multiple pathogenic GoF variants. The convergence of network phenotypes across distinct *KCNH1* variants and their shared response to ASO treatment further suggest that therapeutic development for genetically heterogeneous DEEs can be guided by the underlying disease mechanism rather than individual pathogenic alleles.

*KCNH1* is particularly well suited to an allele-agnostic suppression strategy because its genetic architecture suggests a broad therapeutic window for reducing gene function. Pathogenic *KCNH1* variants are predominantly heterozygous missense variants that increase Kv10.1 channel activity^3–6,14^, whereas population genetic data indicate substantially greater tolerance to LoF than to missense variation^7^. Moreover, direct pharmacological inhibition of Kv10.1 is complicated by its homology to the cardiac potassium channel Kv11.1^5^, making selective suppression of the *KCNH1* transcript an attractive alternative therapeutic approach. Our findings provide experimental evidence for a therapeutic window for such a strategy as approximately 70% reduction of *KCNH1* expression did not measurably perturb spontaneous network activity in control human neurons, yet the same degree of suppression was sufficient to normalize network dysfunction across two distinct pathogenic variants. Similarly, approximately 30% *Kcnh1* suppression was well tolerated *in vivo* and markedly reduced disease severity in *Kcnh1*^Q/+^ mice.

More broadly, our findings suggest a framework for identifying other GoF neurological disorders that may be amenable to allele-agnostic suppression. Candidate genes should combine evidence that pathogenic variants converge on a shared GoF mechanism with evidence that partial LoF is tolerated (Extended Data Fig. 15a). Population genetic constraint analysis (Extended Data Fig. 15b) provides one means of assessing the latter, which can be complemented by phenotypes from genetic knockout animal models and direct experimental evaluation of gene suppression. These criteria may help distinguish GoF disorders in which allele-agnostic suppression offers a scalable therapeutic strategy from those in which preservation of wild-type gene function will require allele-selective approaches.

Our human iPSC neuronal models reveal that distinct *KCNH1* GoF variants converge on a shared network phenotype despite their differential effects on channel kinetics^6^. Both I494V and G375R neurons exhibited a hyperpolarized RMP while largely preserving AP properties yet produced abnormal burst dynamics during neuronal maturation. These findings suggest that increased Kv10.1 activity produces consequences that emerge prominently at the level of developing neuronal networks and highlight the value of human iPSC-derived neuronal MEA recordings for identifying disease phenotypes not readily captured by conventional measures of intrinsic excitability. Importantly, these cellular phenotypes translated to therapeutic efficacy *in vivo* as a single postnatal ICV administration of the mouse-specific ASO substantially reduced seizure burden and improved survival in *Kcnh1*^Q/+^ mice, supporting the feasibility of postnatal gene suppression for severe *KCNH1*-related disease.

Several limitations should be considered. First, although our studies established the therapeutic efficacy of *KCNH1* suppression, the molecular and cellular mechanisms linking altered Kv10.1 activity to network dysfunction were not resolved. Our human neuronal studies focused on excitatory glutamatergic neurons and therefore do not capture the potential contributions of inhibitory neurons or alterations in excitatory–inhibitory circuit balance to *KCNH1*-associated disease. Second, the therapeutic window for *KCNH1* suppression requires further definition, including the degree of reduction that can be tolerated across neuronal populations and developmental stages. Finally, translation will require studies defining the durability, biodistribution, and long-term safety of *KCNH1*-targeting ASOs. Nevertheless, our findings establish that partial *KCNH1* suppression can reverse disease-associated neuronal network dysfunction and ameliorate seizures and early mortality *in vivo*. More broadly, they support a therapeutic framework in which genetically heterogeneous GoF disorders may be treated through allele-agnostic suppression when pathogenic variants converge on a shared mechanism and partial loss of gene function is tolerated.

## Acknowledgments

We are grateful to the following funding sources: US National Institutes of Health (NIH), National Institute on Neurological Disorders and Stroke (NINDS) R21 NS150783 and R01 NS137587 (E.K. & A.L.G), U54 NS108874 (E.K., J.A.K, A.L.G), the New York Stem Cell Foundation (E.K.), the Cure KCNH1 Foundation (E.K., J.A.K), the Karen Toffler Charitable Trust Fellowship (W.C.), and an American Epilepsy Society Predoctoral Fellowship (E.E.C.).

---

**Extended Data Fig. 1.**
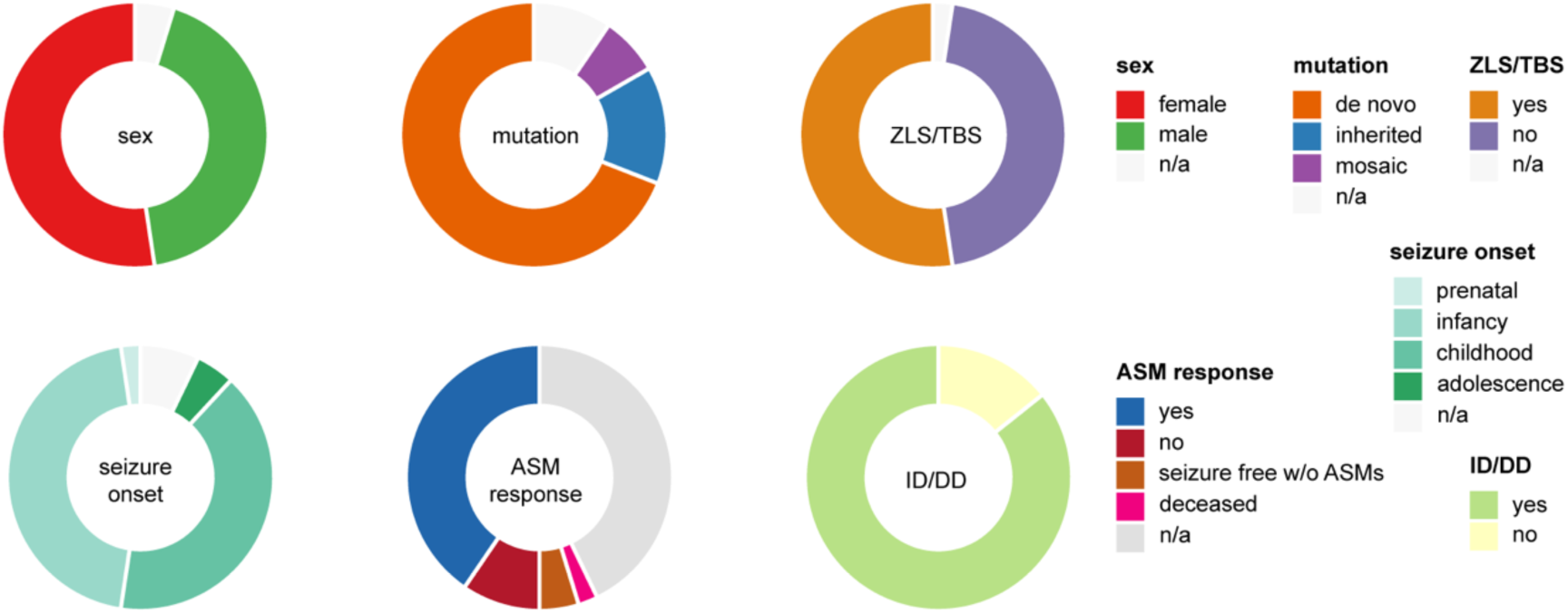
Clinical features of *KCNH1*-related DEEs. ZLS: Zimmermann-Laband syndrome; TBS: Temple-Baraitser syndrome; ASM: antiseizure medication; ID: intellectual disability; DD: developmental delay. Data were adapted from Tian *et al*^14^.

**Extended Data Fig. 2.**
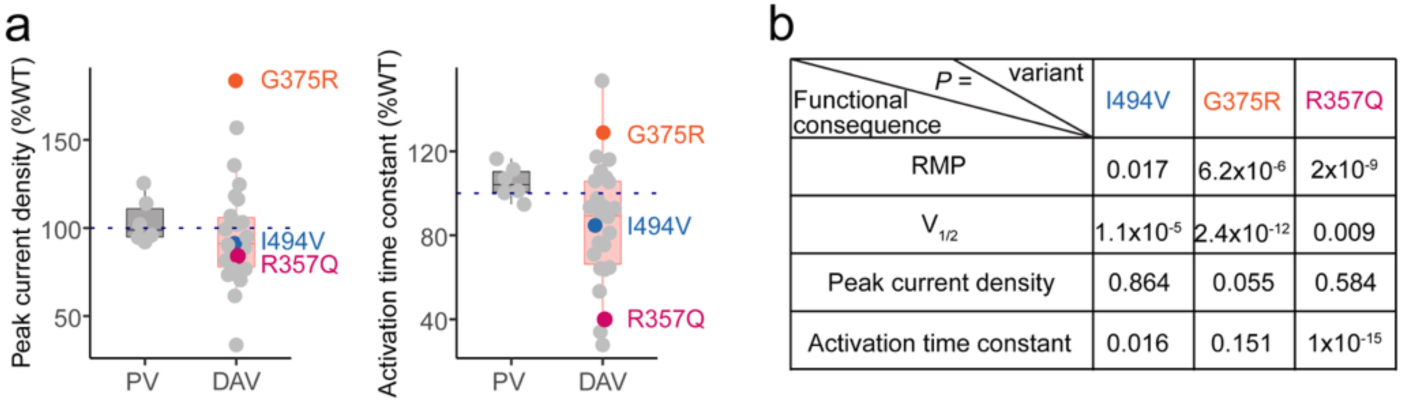
Functional effects of disease-associated variants (DAVs) and population variants (PVs) when expressed in HEK293T cells. **a,** Peak current density and Activation time constant are shown relative to wild-type (WT) channels. **b**, *P*-values of resting membrane potential (RMP), half-activation voltage (V_1/2_), peak current density, and activation time constant in mutant channels in comparison to WT channels. Data were from Thompson *et al*.^6^

**Extended Data Fig. 3.**
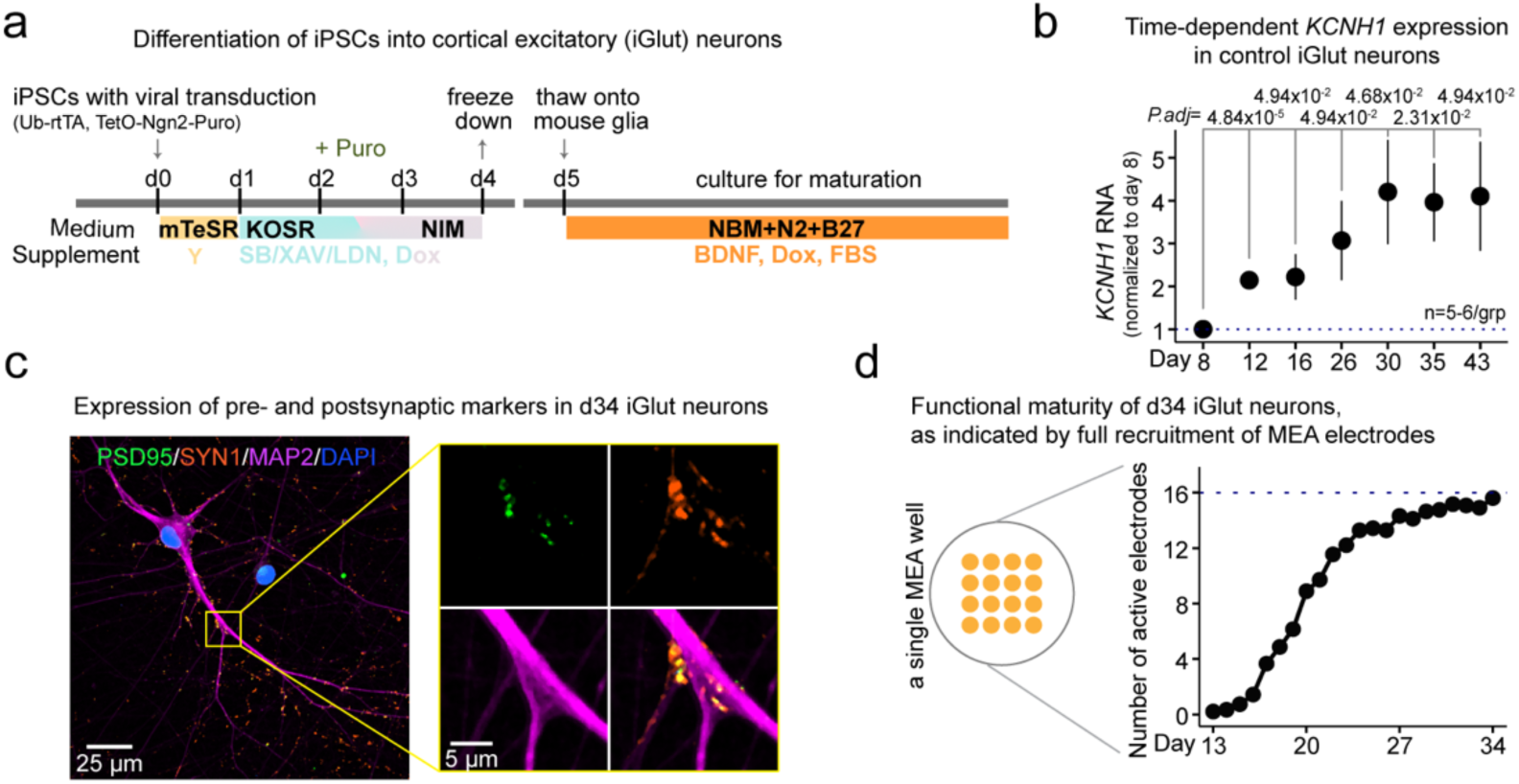
Differentiation and characterization of control iGlut neurons. **a**, Protocol used to differentiate iPSCs into cortical excitatory (iGlut) neurons^23^. **b**, Time-dependent *KCNH1* expression in control iGlut neurons. n=5-6 per condition. Statistical comparisons were performed using one-way ANOVA followed by Tukey’s post hoc test. **c**, Representative expression of the presynaptic marker SYN1 and postsynaptic marker PSD95 in day 34 iGlut neurons. MAP2 and DAPI label neurons and nuclei, respectively. **d**, Number of active electrodes as a function of time during MEA recordings of control iGlut neurons. n=59 wells from six independent differentiation batches.

**Extended Data Fig. 4.**
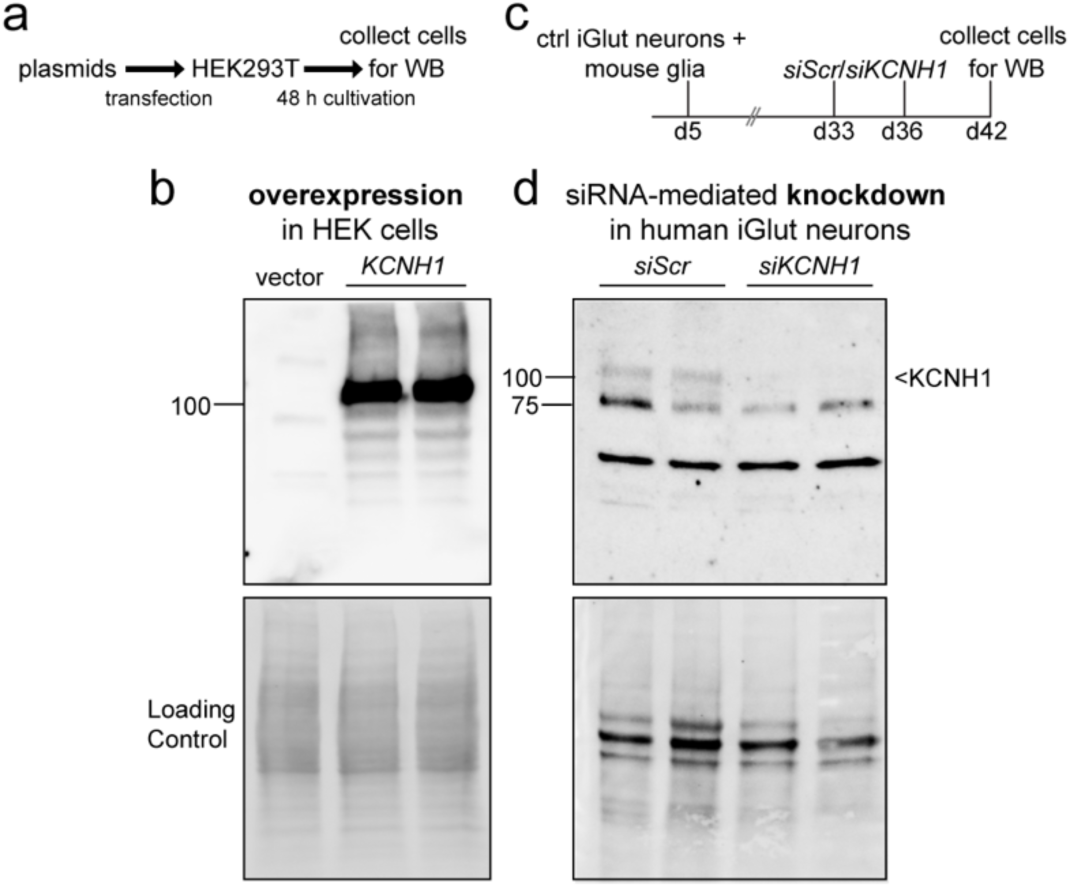
Validation of the *KCNH1* antibody by overexpression and knockdown approaches. **a**, Schematic of the *KCNH1* overexpression experimental workflow. **b**, Western blot analysis of KCNH1 protein following overexpression. **c**, Experimental timeline of siRNA-mediated *KCNH1* knockdown in human iGlut neurons. **d**, Western blot analysis of KCNH1 protein following siRNA-mediated knockdown.

**Extended Data Fig. 5.**
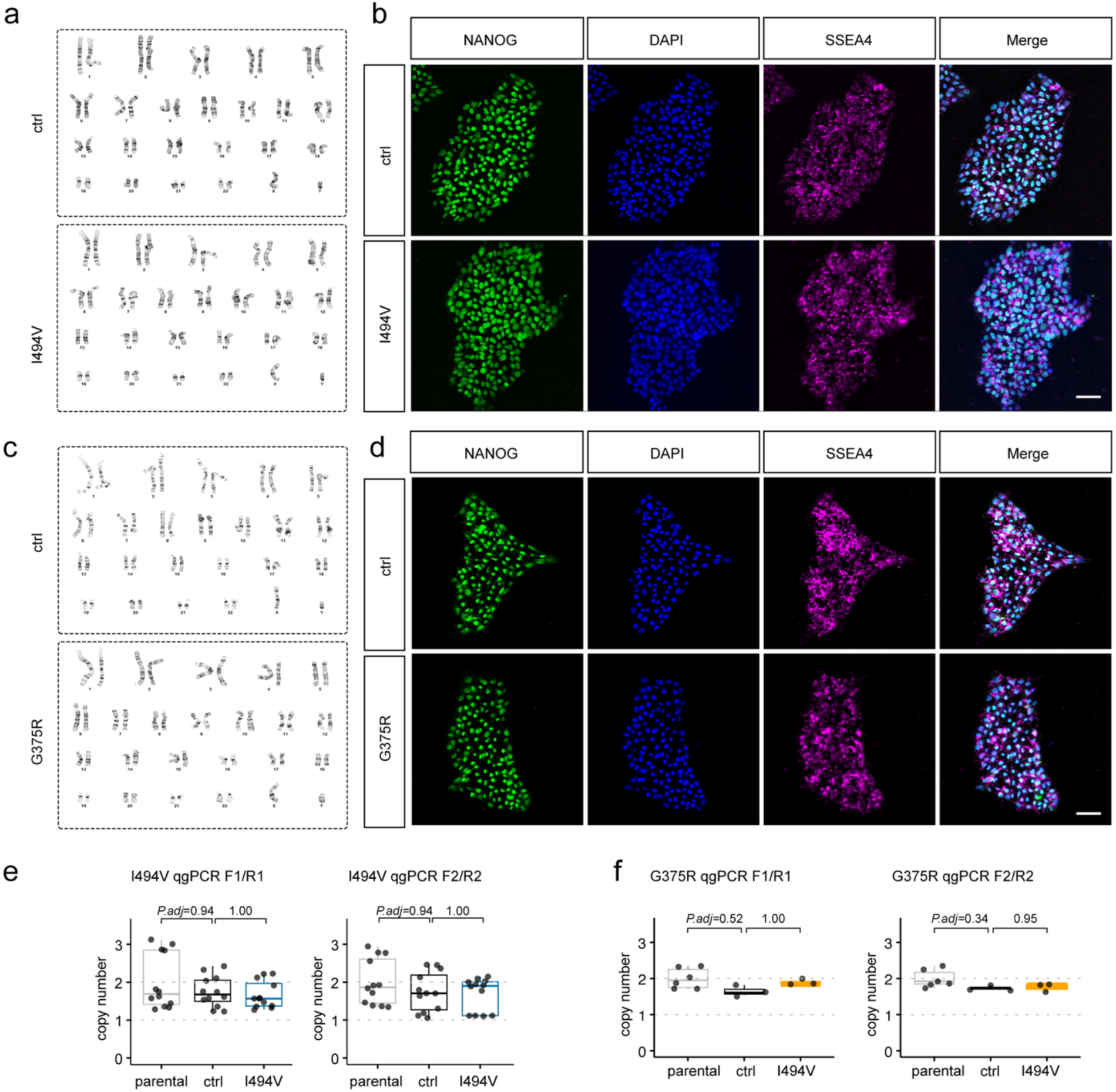
Control and *KCNH1* knock-in iPSC lines exhibit normal karyotypes, pluripotency, and *KCNH1* copy number. **a**,**c**, G-banded karyotypes of control and mutant iPSC lines. **b**,**d**, Immunostaining for the pluripotency markers NANOG and SSEA4 in control and mutant iPSC lines. DAPI labels nuclei. Scale bar, 50 μm. **e**,**f**, Quantitative genomic PCR confirms comparable *KCNH1* copy number in control and mutant iPSC lines. n=12 samples per I494V condition and n=3-6 samples per G375R condition. Statistical comparisons were performed using one-way ANOVA followed by Tukey’s post hoc test.

**Extended Data Fig. 6.**
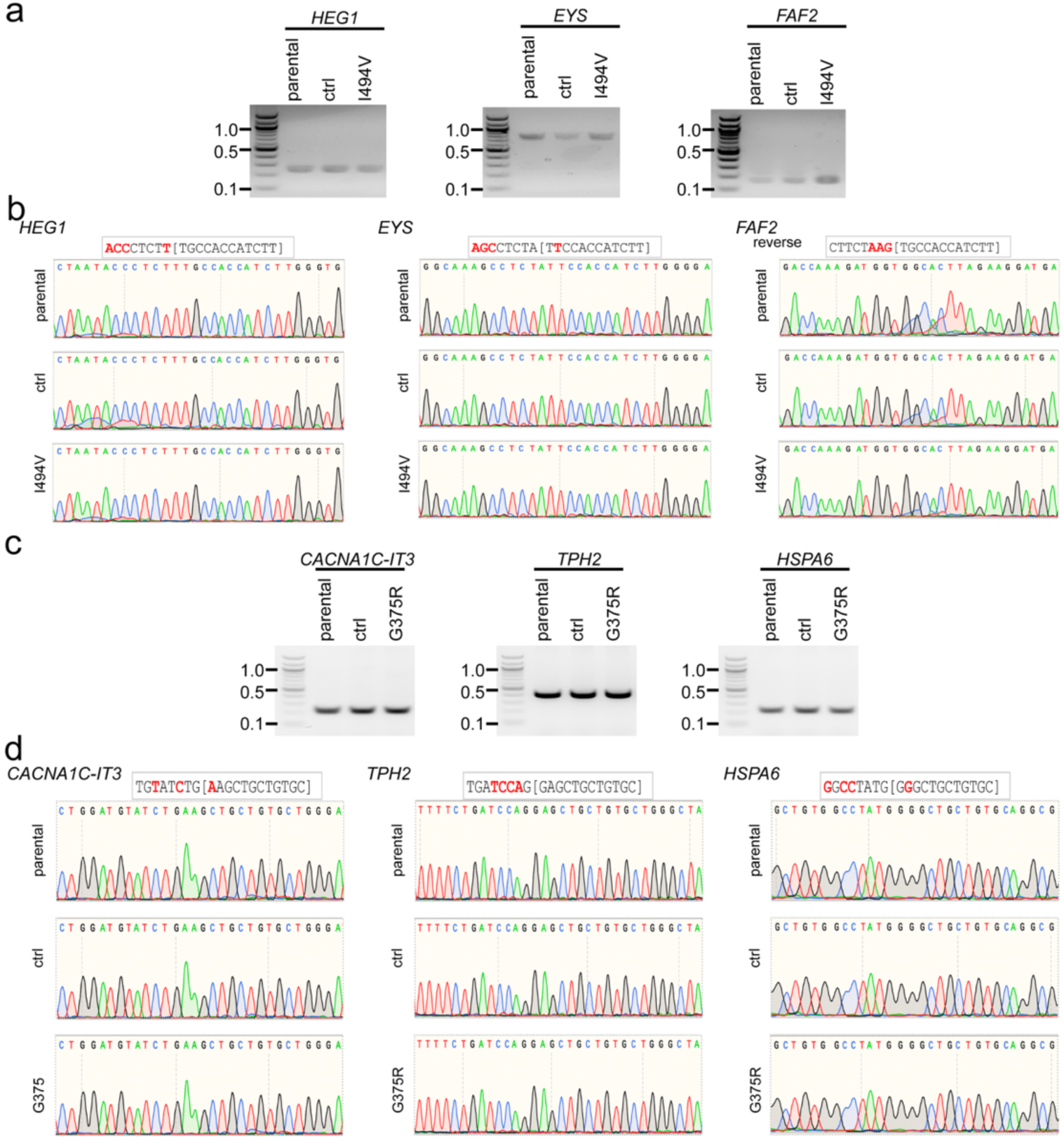
No detectable mutations at predicted CRISPR off-target sites by PCR and Sanger sequencing. **a**, Agarose gel electrophoresis of PCR products spanning the predicted off-target loci *HEG1*, *EYS*, and *FAF2* amplified from the parental iPSC line, edited control (ctrl), and I494V. **b**, Sanger sequencing chromatograms of the PCR products shown in a, confirming the absence of sequence alterations at the predicted off-target sites. **c**, Agarose gel electrophoresis of PCR products spanning the predicted off-target loci *CACNA1C-IT3*, *TPH2*, and *HSPA6* amplified from the parental iPSC line, ctrl, and G375R. **d**, Sanger sequencing chromatograms of the PCR products shown in c, confirming the absence of sequence alterations at the predicted off-target sites.

**Extended Data Fig. 7.**
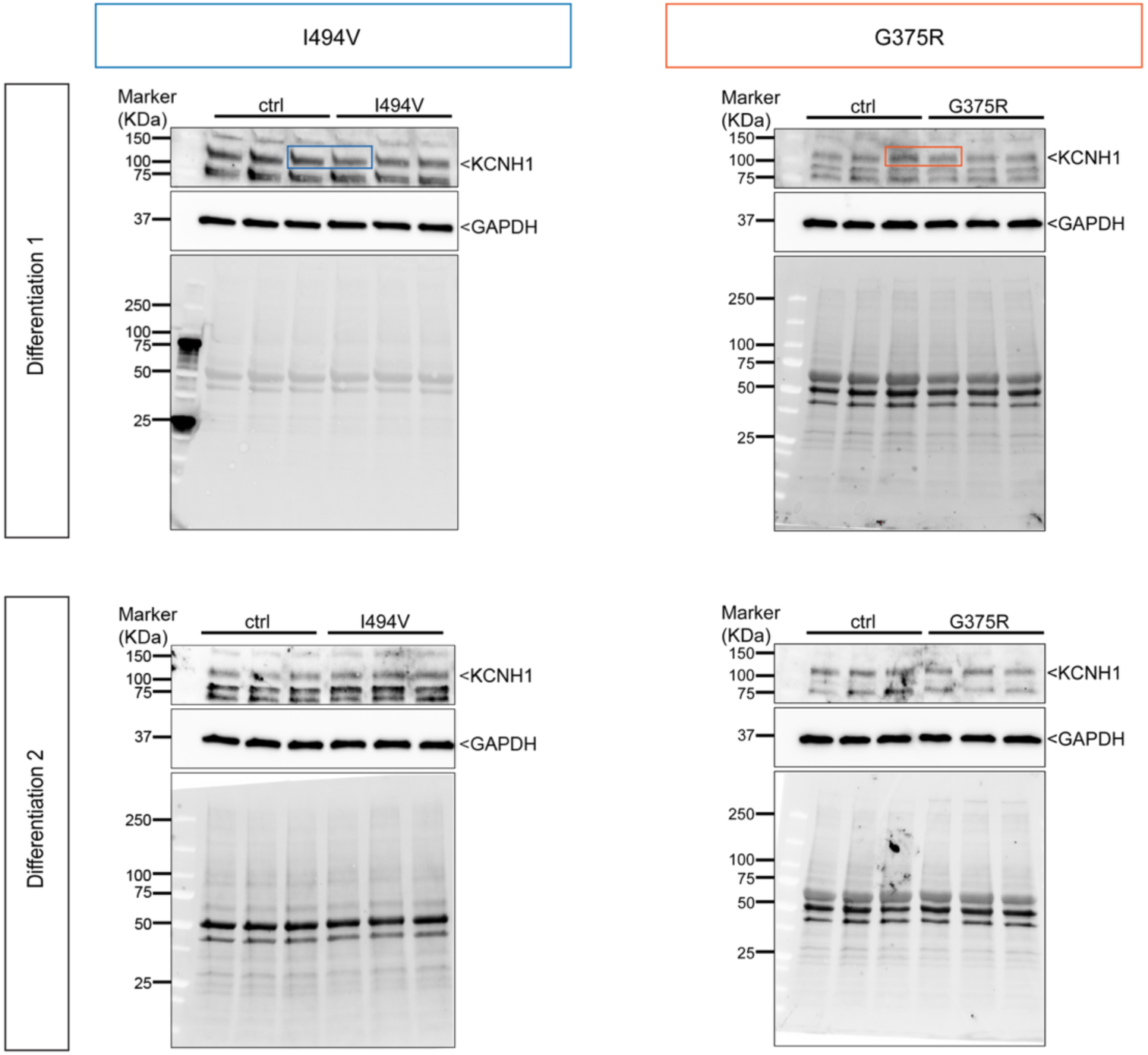
Western blot analysis of KCNH1 protein expression in control and mutant iGlut neurons. Immunoblots showing KCNH1 protein expression in control, I494V, and G375R iGlut neurons from two independent differentiations. Quantification of KCNH1 protein levels normalized to GAPDH is shown in Fig. 2c.

**Extended Data Fig. 8.**
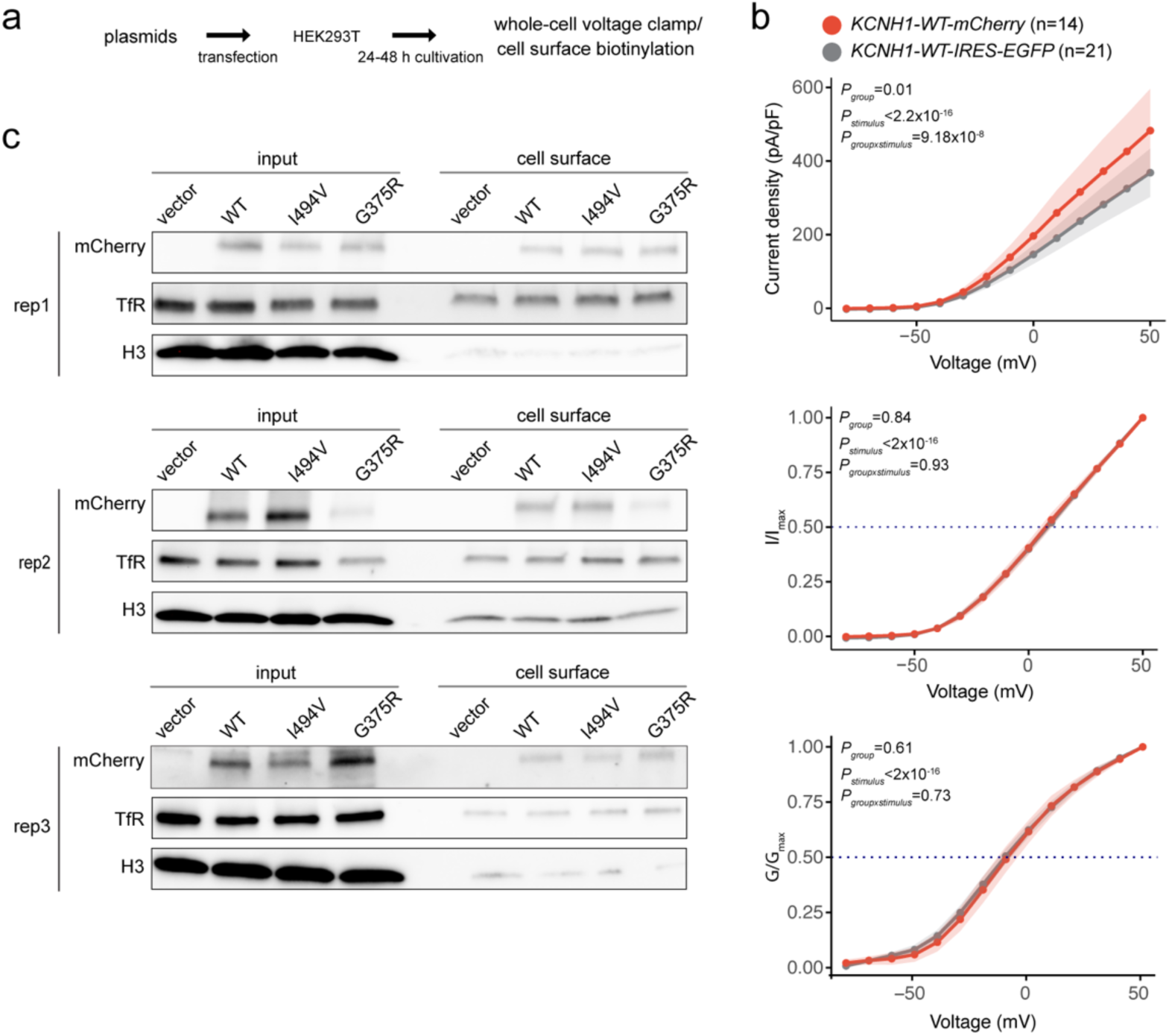
I494V and G375R mutations do not impair Kv10.1 protein trafficking in a heterologous expression system. **a,** Schematic of the experimental workflow and timeline. **b**, C-terminal mCherry tagging slightly increased the steady-state current density but did not affect normalized current and normalized conductance. n=14-21 cells per group. Data were presented mean±95% confidence interval and were analyzed using a linear mixed-effects model with group, stimulus intensity, and their interaction as fixed effects, and cell identity as a random intercept to account for repeated measurements from the same cell. Fixed effects were evaluated using Type III F-tests with Satterthwaite’s approximation for denominator degrees of freedom. **c**, Western blot analysis of total input and cell surface protein fractions from cells expressing empty vector, WT-mCherry, I494V-mCherry, or G375R-mCherry. n=3 independent transfection. TfR, transferrin receptor, a marker of the plasma membrane fraction; H3, histone H3, a nuclear protein marker. Quantification of the ratio of cell-surface to total KCNH1 protein is shown in Fig. 2d.

**Extended Data Fig. 9.**
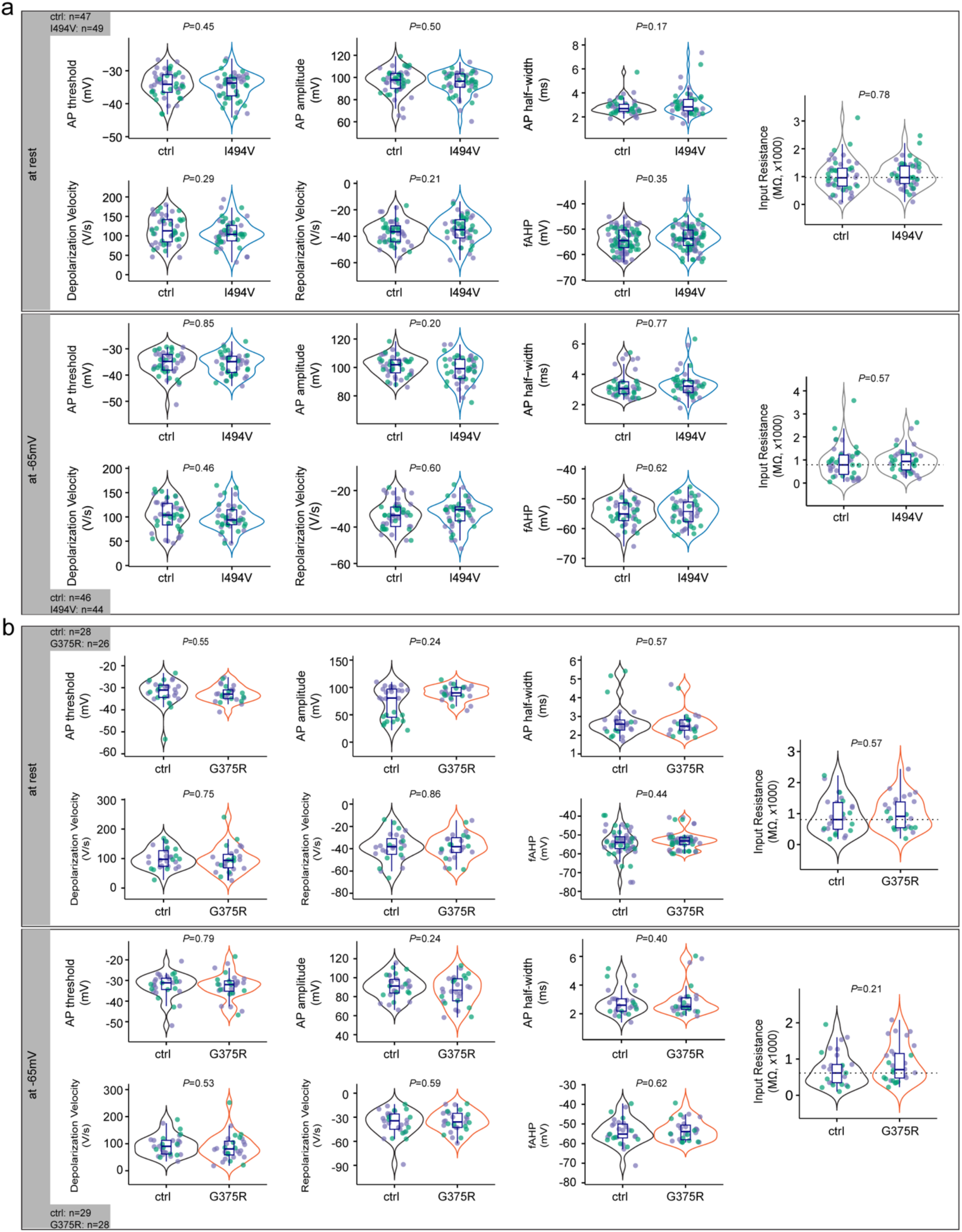
I494V and G375R mutations do not alter action potential (AP) properties or input resistance. **a,** I494V mutant neurons exhibit comparable input resistance and action potential properties, including threshold, amplitude, half-width, depolarization velocity, repolarization velocity, and fast afterhyperpolarization (fAHP), relative to control neurons. Measurements were performed at the resting membrane potential or after adjusting cells to −65 mV. n=44-49 neurons per condition from two independent differentiation batches. Statistical comparisons were performed using Welch’s two-sample *t*-test**. b,** G375R mutant neurons similarly exhibit preserved action potential properties. n=26-29 neurons per condition from two independent differentiation batches. Statistical comparisons were performed using Welch’s two-sample *t*-test.

**Extended Data Fig. 10.**
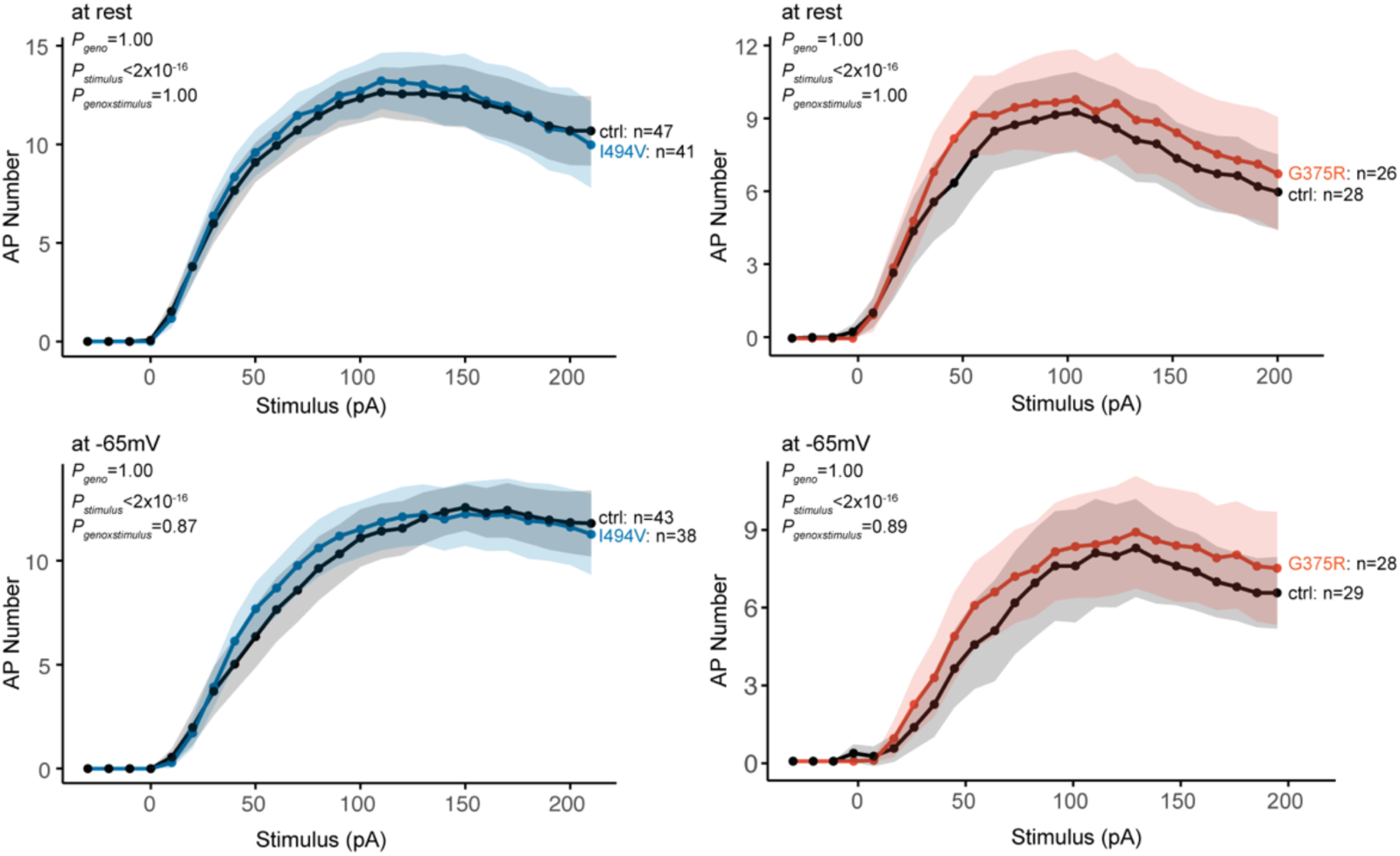
Evoked action potential number is comparable between control and mutant neurons. Measurements were performed at the resting membrane potential or after adjusting cells to −65 mV. n=38-47 neurons per I494V condition from two independent differentiation batches and n=26-29 neurons per G375R condition from two independent differentiation batches. Data were presented mean±95% confidence interval and were analyzed using a generalized linear mixed-effects model with genotype, stimulus intensity, and their interaction as fixed effects, and cell identity as a random intercept to account for repeated measurements from the same cell. Overall effects were evaluated using Type III Wald χ² tests.

**Extended Data Fig. 11.**
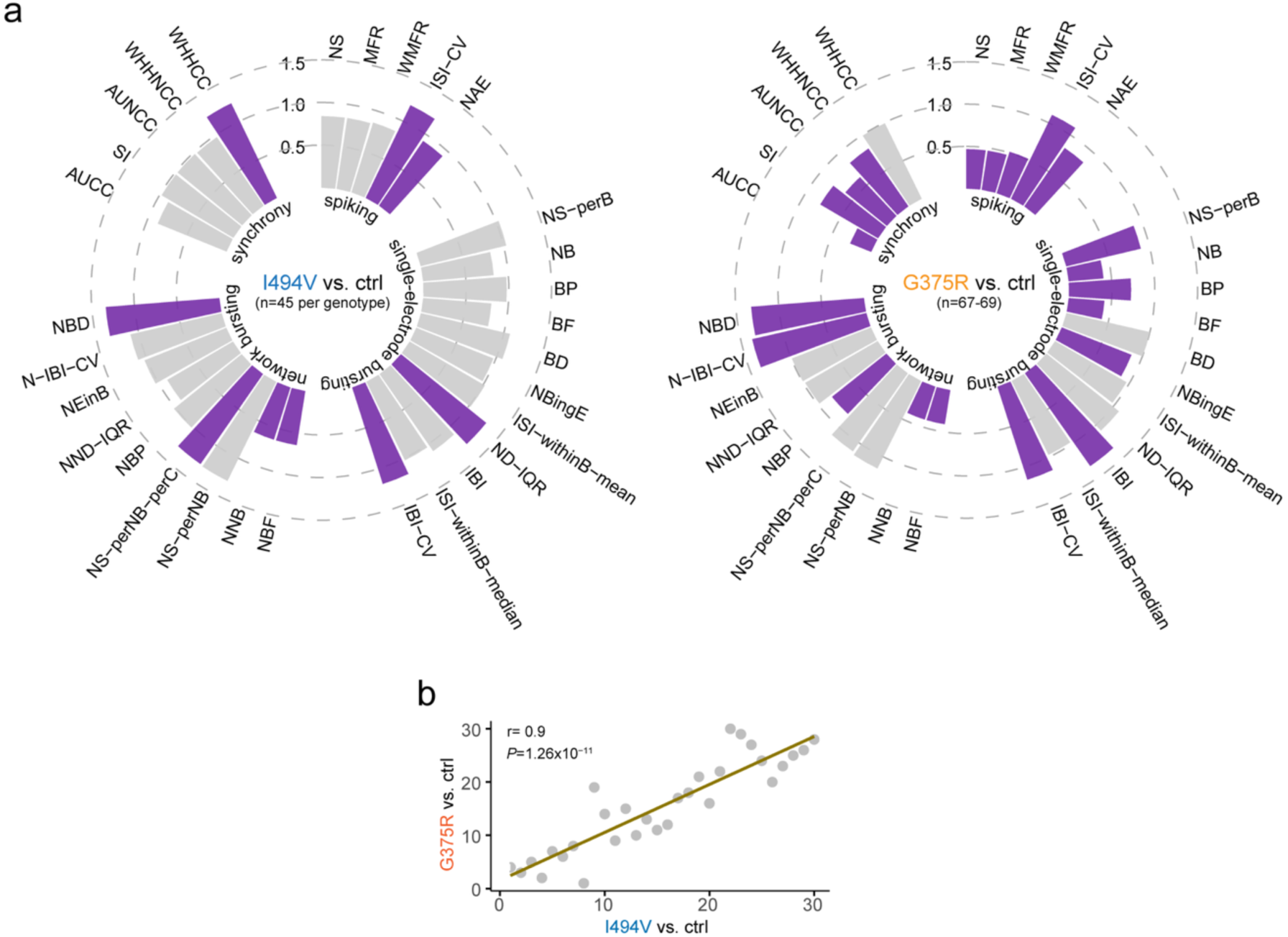
MEA phenotypes from I494V and G375R mutant neurons across days 13-32. **a**, Circular bar plot showing area under the curve (AUC) comparisons of 30 MEA parameters measured from days 13-32 in control and I494V or G375R mutant neuronal lines. n=45-69 wells per condition from three to four independent differentiation batches. Statistical comparisons were performed using Wilcoxon rank-sum test, followed by Benjamini-Hochberg correction for multiple comparisons across 30 parameters. **b,** Correlation analysis of ranked MEA parameters between I494V and G375R mutant neurons. Statistical significance was assessed using Spearman’s rank correlation test.

**Extended Data Fig. 12.**
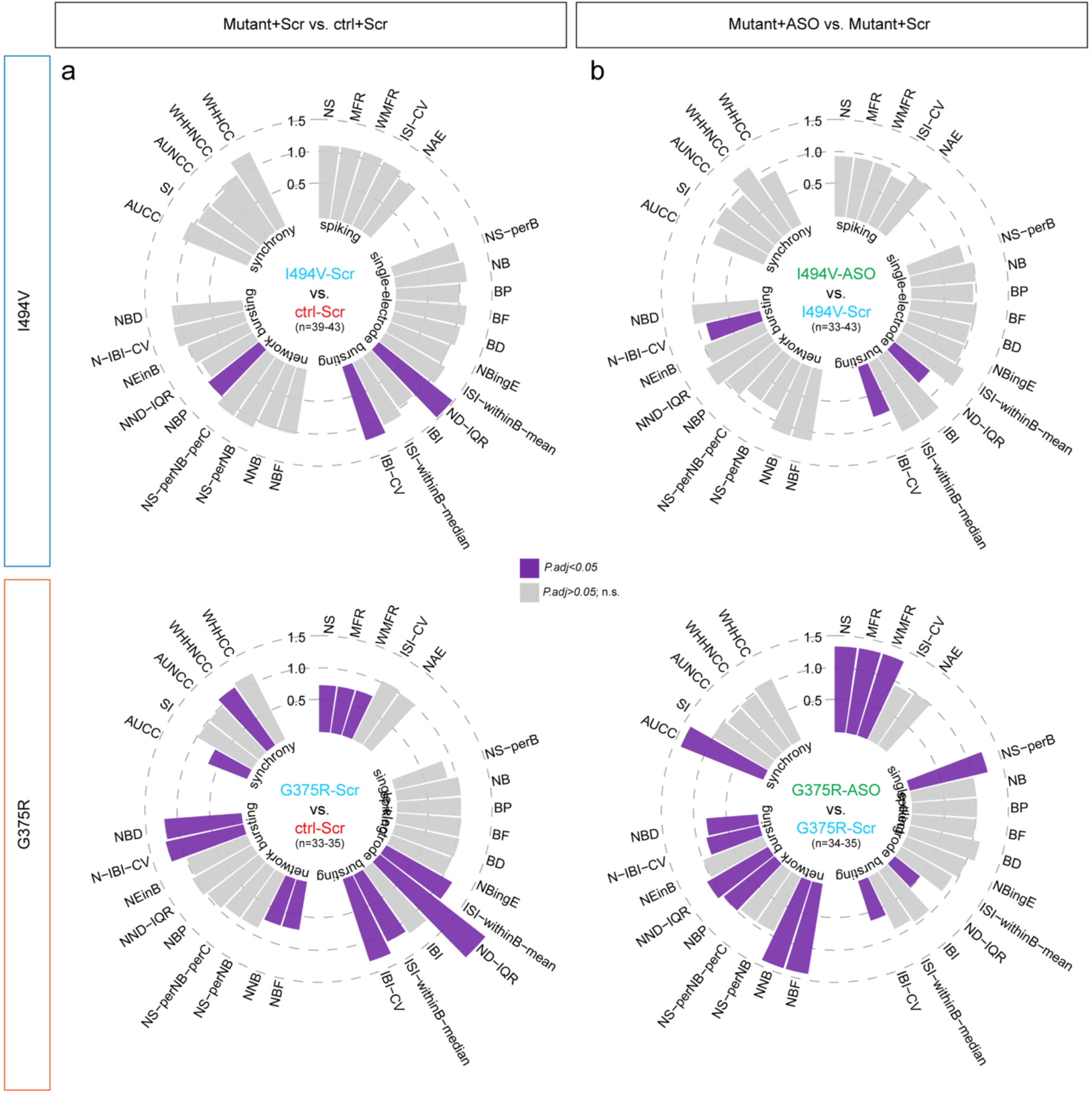
Antisense suppression of *KCNH1* reverses neuronal network dysfunction caused by I494V or G375R. MEA parameters from mutant neurons were normalized to those from their corresponding controls. n=33-43 wells per condition from three to four independent differentiation batches. For each MEA parameter, genotype difference (**a**), treatment effect (**b**), or rescue effect (Fig. 3g-h) was assessed using the Kruskal-Wallis test, followed by Benjamini-Hochberg correction across the 30 parameters.

**Extended Data Fig. 13.**
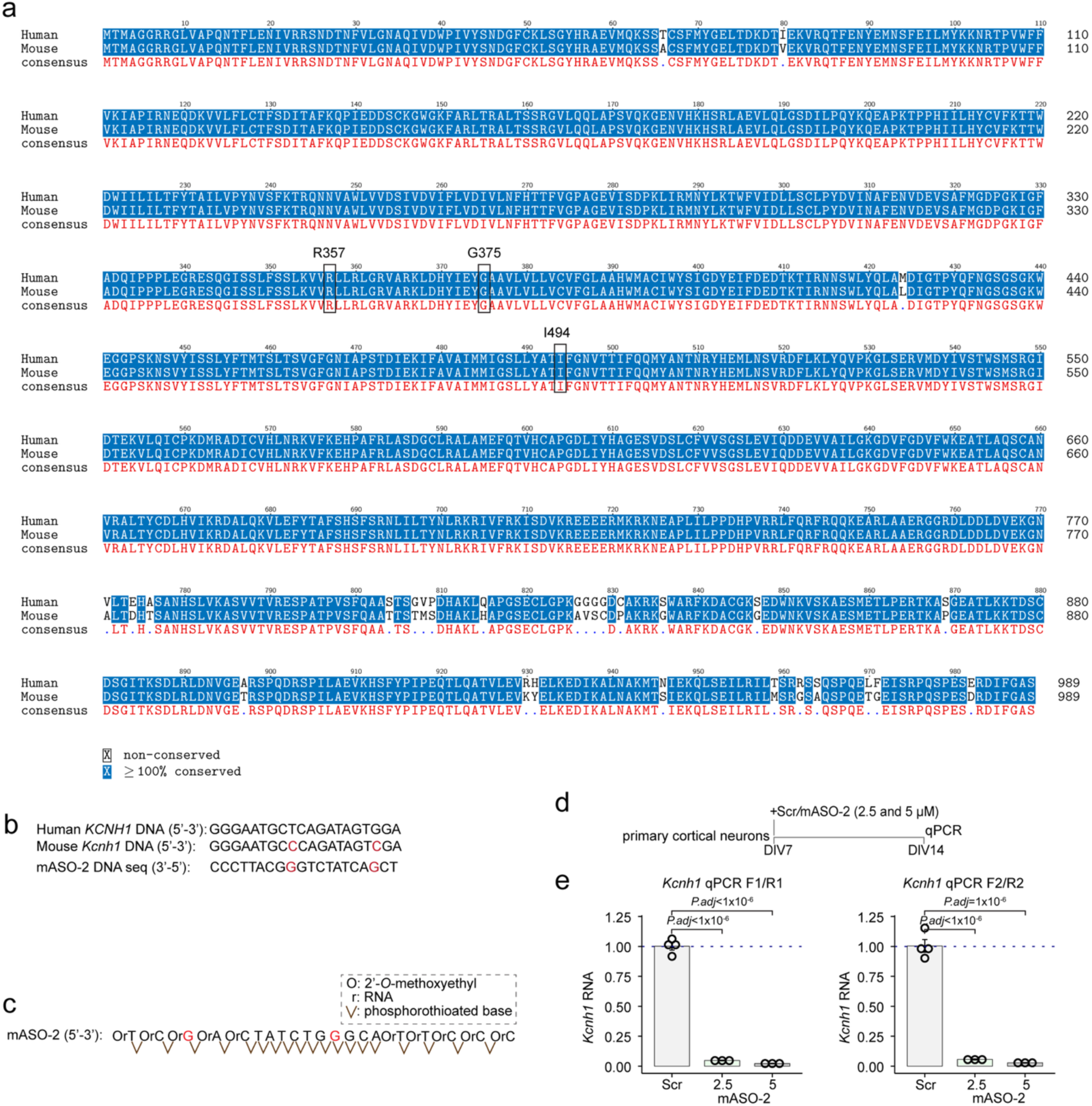
Design and validation of mouse *Kcnh1*-targeting ASOs. **a**, Alignment of human and mouse KCNH1 protein sequences. The three mutations examined in this study are labelled and are conserved between human and mouse. **b**, Mouse *Kcnh1* ASO DNA sequence and corresponding genomic sequences in human and mouse. **c**, Sequences and chemical modifications of the mouse *Kcnh1*-targeting ASO (mASO-2). **d**, Experimental timeline for testing *Kcnh1*-targeting ASOs in primary mouse cortical neurons. **e,** *Kcnh1* expression following treatment with scrambled (Scr) or *Kcnh1*-targeting ASOs. Data were normalized to the house-keeping genes *Gapdh* and *Pgk1*. n=3-4 wells per condition. Statistical comparisons were performed using one-way ANOVA followed by Tukey’s post hoc test.

**Extended Data Fig. 14.**
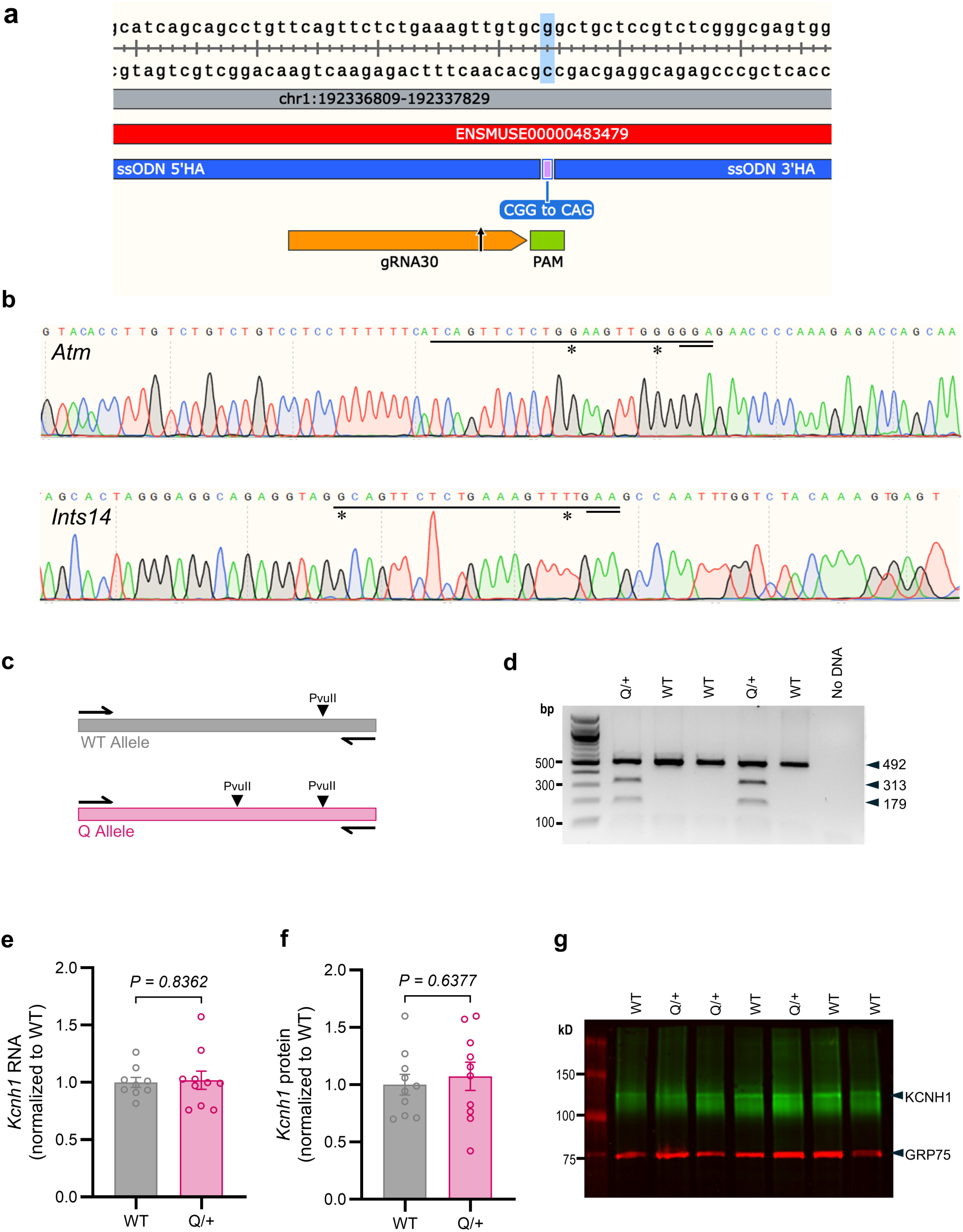
Generation of *Kcnh1^R357Q^*mouse line. **a**, Schematic of CRISPR/Cas9 targeting strategy. **b**, Chromatograms from Sanger sequencing of amplicons for predicted intergenic off-targets with <3 mismatches confirmed the absence of sequence alterations at the predicted off-target sites. **c**, Schematic of restriction length polymorphism (RFLP)-based genotyping of *Kcnh1^Q/+^* mice by PvuII digestion of 542 bp PCR amplicon with primers *Kcnh1*-R357Q forward and *Kcnh1*-R357Q reverse (See Supplemental Table 4 for primer sequences). **d,** Representative RFLP results following PvuII digestion showing predicted 492-bp fragment for wild-type (WT) and 492-, 313-, and 179-bp fragments for *Kcnh1^Q/+^* (Q/+) heterozygotes. Both alleles also generate a 50-bp band that is not shown. **e**, Brain expression of *Kcnh1* transcript did not differ between *Kcnh1^Q/+^* (Q/+, n=9) and wild-type (WT, n=10) by RT-ddPCR. **f**, Forebrain expression of Kv10.1 protein did not differ between *Kcnh1^Q/+^* (Q/+, n=10) and wild type (WT, n=10) by western blotting. **g**, Representative western blot of forebrain membrane proteins (50 µg) probed for KCNH1 protein and GRP75 as a normalization control. Treatment groups were compared using Welch’s two-sample *t*-test and each circle represents an individual mouse in panels e and f.

**Extended Data Fig. 15.**
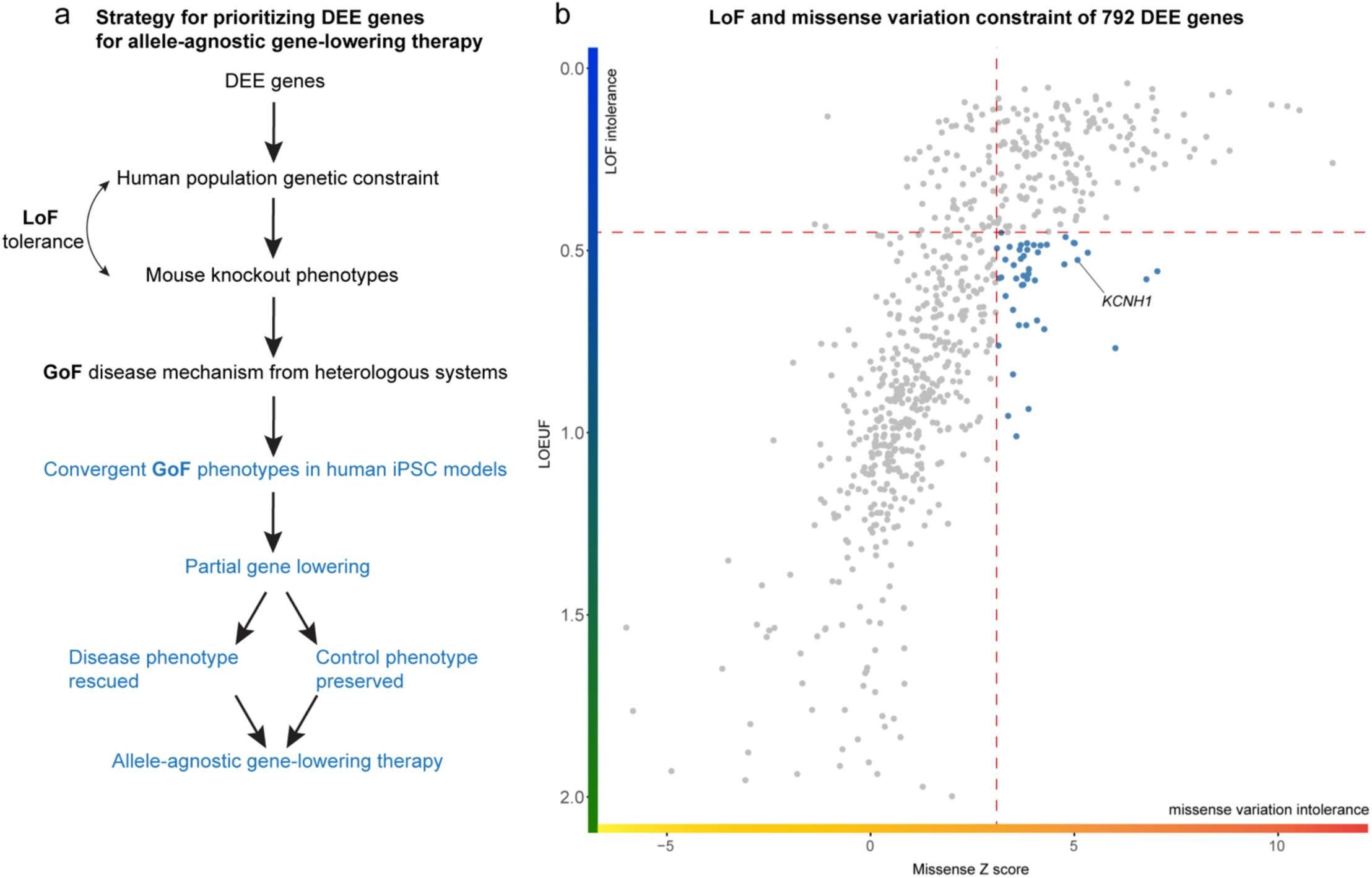
Strategy for identifying DEE genes potentially amenable to allele-agnostic gene-lowering therapy. **a**, Candidate DEE genes are prioritized based on genetic and functional evidence and modeled in human iPSC-derived neurons. Disease-relevant neuronal phenotypes are quantified and compared between mutant and control lines. Genes for which pathogenic variants produced a convergent disease phenotype that can be reversed by partial gene reduction are considered candidates for allele-agnostic gene-lowering therapy. **b**, Gene-level constraint metrics (gnomAD^7^, v4.1) are used to prioritize candidates. Each dot indicates one gene and the DEE gene list was obtained from Oliver *et al.*^10^ (v2026-03). Genes in blue represent those meeting the selected constraint criteria, with dashed lines indicating the applied thresholds. LOEUF: loss-of-function observed/expected upper bound fraction.

**Extended Data Video 1.** Representative tonic-clonic seizure in a *Kcnh1^Q/+^* mouse chronically treated with GS967. Tonic-clonic seizures in GS967-treated *Kcnh1^Q/+^* mice typically terminate without progressing to tonic hindlimb extension.

**Extended Data Video 2.** Rare tonic seizure with hindlimb extension in a *Kcnh1^Q/+^*mouse treated with mASO-2 (30 µg) at P1.

**Extended Data Video 3.** Representative tonic seizure with hindlimb extension in a *Kcnh1^Q/+^* mouse treated with vehicle at P1.

**Extended Data Video 4.** Representative tonic seizure with hindlimb extension observed in a *Kcnh1^Q/+^* mouse not treated with mASO-2 or vehicle.

## Methods

### Cell culture

Human induced pluripotent stem cells (iPSCs) were maintained on Matrigel-coated plates in mTeSR media (STEMCELL Technologies) and passaged twice weekly using Accutase (Sigma-Aldrich). Primary mouse glial cells were maintained in Dulbecco’s Modified Eagle Medium (DMEM) supplemented with 10% fetal bovine serum and GlutaMAX. Primary mouse cortical neurons were maintained in Neurobasal-A medium supplemented with B27, GlutaMAX, and CultureOne (Thermo Fisher). Induced glutamatergic (iGlut) neurons were co-cultured with primary mouse glial cells. Briefly, glial cells were plated onto poly-D-lysine/laminin -coated culture plates or coverslips in glial media consisting of Minimum Essential Medium (MEM; Thermofisher) supplemented with GlutaMAX, D-glucose, and horse serum. Three days later, frozen iGlut neurons were thawed and plated onto the glial monolayer in complete Neurobasal medium consisting of Neurobasal media supplemented with non-essential amino acids (NEAA), GlutaMAX, N2, B27, 10 ng/mL brain-derived neurotrophic factor (BDNF; R&D Systems), 2% fetal bovine serum (FBS), 3 μg/mL doxycycline, and 10 μM Y-27632 Rock inhibitor. Cultures were maintained in complete Neurobasal medium with half-medium changes three times per week. All cell culture were maintained at 37°C in a humidified atmosphere containing 5% CO2 without antibiotics and were routinely confirmed to be free of mycoplasma contamination (MycoAlert Mycoplasma Detection Kit).

### Primary mouse glial preparation

Primary glial cells were prepared from the cerebral cortices of postnatal day 0-1 CD-1 mice (Charles River Laboratories). Briefly, cortices were dissected in Hank’s Balanced Salt Solution (HBSS, Ca^2+^ and Mg^2+^ free; Thermo Fisher), enzymatically dissociated with trypsin (Thermo Fisher) and DNaseI (Worthington Biochemical Co.), and passed through a 40 μm cell strainer (Fisher Scientific) to a single-cell suspension. Cells were plated and cultured in MEM supplemented with 10% horse serum, 6% D-glucose, 1% GlutaMAX, and 1% penicillin-streptomycin. After 24 h, the medium was replaced with the same formulation without antibiotics, and cultures were maintained until confluent. Confluent glial cultures were passaged once at a 1:3.5 split ratio, expanded, and cryopreserved for subsequent co-culture experiments.

### Primary mouse cortical neuron preparation

Primary cortical neurons were prepared from mouse embryos. Cortices were dissected in dissection medium consisting of HBSS (Ca^2+^ and Mg^2+^ free) supplemented with sodium pyruvate, 0.1% glucose and 10 mM HEPES. Tissue was enzymatically dissociated with papain in the presence of DNase I to minimize cell clumping, followed by gentle trituration to generate a single-cell suspension. Cells were plated in Basal Medium Eagle (BME) supplemented with 10% FBS, 0.45% glucose, sodium pyruvate, GlutaMAX, and penicillin-streptomycin. Following cell attachment, cultures were maintained in Neurobasal-A medium supplemented with B27, GlutaMAX, and CultureOne. Half of the culture medium was replaced once per week.

### Generation of knock-in iPSCs

The parental iPSC line KOLF2.1J was obtained from John Landers (University of Massachusetts) and has been extensively characterized previously^24^. *KCNH1* knock-in (KI) lines carrying the I494V or G375R variants were generated from KOLF2.1J by CRISPR-Cas9-mediated genome editing using guide 5’-CTTCTCTATGCCACCATCTT or 5’-TGAATATGGAGCTGCTGTGC, respectively (EditCo). All edited lines were validated by Sanger sequencing and confirmed to have normal karyotypes, intact pluripotency, and normal *KCNH1* copy number.

### Lentiviral production and transduction

Lentiviruses used in this study, including FUW-M2rtTA (Addgene #20342)^25^, TetO-Ngn2-Puro (Addgene #52047)^26^, and CamKIIA-EGFP (Addgene #112670)^27^, were prepared by GET iN Core at Northwestern University, which is supported by NIH under award number P30AR075049. For transduction, viruses were incubated with iPSCs or neurons for 5 min at room temperature and cells were then plated.

### Plasmid construction, preparation, and transfection

Expression constructs encoding wild-type or mutant *KCNH1* (NM_002238) were generated commercially (VectorBuilder). *KCNH1* was fused with in-frame to a C-terminal mCherry tag and expressed under the control of the CMV promoter. Plasmid sequences were verified by whole-plasmid sequencing (Plasmidsaurus), and plasmids were purified using the ZymoPUREII Plasmid Purification Kit (Zymo Research) according to the manufacturer’s instructions. Transfection was done using Lipofectamine 3000 Transfection Reagent (Thermo Fisher) according to manufacturer’s instructions.

### siRNA-mediated *KCNH1* knockdown

The siRNAs were from Thermo Fisher (siScr: 4390844, siKCNH1: s7730, s7731). Co-cultured cells were treated twice with siRNAs on day 33 and day 36 using Lipofectamine RNAimax Transfection Reagent (ThermoFisher) and final siRNA concentration was 60 nM. Cells were collected for WB on day 42 (Extended Data Fig. 3c-d).

### Human iGlut neuron differentiation

Differentiation into iGlut neurons was performed using an established NGN2-based protocol (Extended Data Fig. 2a)^23^. Briefly, iPSCs were dissociated into single cells using Accutase and transduced with lentiviruses encoding FUW-M2rtTA and TetO-Ngn2-Puro. Neuronal differentiation was induced by doxycycline and dual-SMAD inhibition using SB431542 and LDN-193189 together with WNT inhibition by XAV939. Puromycin selection was applied to enrich transduced cells. Over the 4-day induction protocol, cultures were transitioned from knockout serum replacement medium containing knockout DMEM, 15% knockout replacement serum, 1% NEAA, 1% GlutaMAX, and 0.1% beta-mecaptoethanol to neural induction medium containing DMEM/F12, 1% N2 supplement, 1% NEAA, 1% GlutaMAX, 0.16% D-glucose, and 0.004% heparin. On day 4, induced neurons were harvested and cryopreserved in 10% dimethyl sulfoxide (DMSO) in FBS for subsequent co-culture with primary mouse glia.

### DNA, RNA, and protein extraction from cells

Genomic DNA (gDNA) and total RNA were extracted using the DNeasy Blood & Tissue Kit (Qiagen) and the NucleoSpin RNA kit (Macherey-Nagel), respectively, according to the manufacturer’s instructions. Complementary DNA (cDNA) was synthesized from total RNA using the iScript Reverse Transcription Supermix (Bio-Rad). For protein extraction, cells were harvested and lysed in NTE buffer (150 mM NaCl, 50 mM Tris-HCl, 5 mM EDTA, 1% SDS) supplemented with a protease inhibitor cocktail (MilliporeSigma). Lysates were sonicated (3 x 30 s) and centrifugated at 15,000 x g for 15 min at 4°C. The clarified supernatant was collected, and protein concentration was determined using Pierce BCA Protein Assay Kit (Thermo Fisher Scientific) before subsequent immunoblotting.

### Genomic integrity, off-target, and *KCNH1* allele copy number analysis

Genomic integrity of all control and mutant iPSC clones were accessed by G-banded karyotyping performed by Cell Line Genetics. Potential CRISPR-Cas9 off-target sites in control and mutant iPSC clones were predicted using the CCTop online tool (Supplemental Table S3)^28^. Three predicted off-target loci were randomly selected for validation using PCR and Sanger sequencing (Northwestern Sanger Sequencing Facility). *KCNH1* copy number was determined by genomic quantitative PCR.

### Quantitative PCR (qPCR)

qPCR was performed in 20 μL reactions containing 25-100 ng of gDNA or cDNA, 0.4 μM of each primer, and 1x Power SYBR Green PCR Master Mix (Thermo Fisher Scientific). Reactions were run on a CFX384 Touch Real-Time PCR Detection System (Bio-Rad) using the following conditions: 50 °C for 2 min; 95 °C for 2 min; 40 cycles of 95 °C for 10 s and 60 °C for 35 s; followed by a dissociation curve analysis at 95 °C for 15s, 65 °C for 5s, and 95 °C for 15 s with a ramp rate of 0.5 °C. Gene expression was quantified using the comparative Ct method and normalized to the housekeeping genes *GAPDH* and *GPI*.

### Immunocytochemistry

Cells cultured on coverslips were fixed with 4% paraformaldehyde (Electron Microscopy Sciences) in phosphate-buffered saline (PBS) for 10 min at room temperature, washed three times with PBS, permeabilized and blocked in 10% normal donkey serum (NDS, Jackson Immuno Research) containing 0.1% Triton X-100 in PBS (PBST) for 1 h at room temperature. Primary antibodies were diluted in PBST containing 2% NDS and incubated overnight at 4°C. After washing, fluorescence-conjugated secondary antibodies (Thermo Fisher Scientific) were applied at a 1:300-1:500 dilution for 1 h at room temperature. Coverslips were mounted with Fluromount-G (SounthernBiotech), and images were acquired using a Nikon CSU-W1 SoRa spinning-disk confocal microscope in the Center for Advanced Microscopy/Nikon Imaging Center at Northwestern University, which is supported by the S10 award (1S10OD032270-01A1). Confocal Z-stack images were converted to maximum-intensity projections using NIS-elements AR (v5.41.00; Nikon). The resulting images were then assembled into montages using FIJI (v2.14.0/1.54f).

### Cell surface protein biotinylation

Cell surface proteins were isolated using the Pierce Cell Surface Protein Isolation Kit (Thermo Fisher) according to the manufacturer’s instructions with minor modifications. Briefly, cells were washed twice with ice-cold PBS and incubated with Sulfo-NHS-SS-Biotin labelling solution for 30 min at 4 °C. Excess biotin was quenched with supplied quenching solution for 15 min at 4 °C. Cells were then gently scraped, collected by centrifugation (500 x g, 3 min, 4 °C), and washed twice with ice-cold Tris-buffered saline (TBS). Cell pellets were lysed in 500 μL of the supplied lysis buffer supplemented with protease inhibitor cocktails and incubated on ice for 30 min. To improve membrane protein solubilization, lysates were sonicated once (10% amplitude, 5 x 1 s pulses) and vortexed briefly every 5 min during lysis. Lysates were clarified by centrifugation (10,000 x g, 2 min, at 4 °C). Of the resulting supernatant, 400 μL was incubated with immobilized NeutrAvidin resin for 1 h at room temperature with end-over-end mixing to capture biotinylated surface proteins, and ∼100 μL was retained as the total input fraction. Following binding, the NeutrAvidin resin was washed three times with wash buffer, and bound proteins were eluted by incubation in 1 x SDS sample buffer (80 mM Tris, 2% SDS, 10% glycerol, 50 mM DTT) for 5 min at 95 °C. Eluted surface fractions and total input samples were analyzed by SDS-PAGE and immunoblotting.

### SDS-PAGE and immunoblotting

Protein lysates were mixed with Laemmli sample buffer (Bio-Rad), denatured at 95°C for 5 min, and separated by SDS-PAGE using Mini-PROTEAN TGX precast gels (Bio-Rad). Proteins were transferred onto nitrocellulose membranes (Bio-Rad), which were then blocked in 5% non-fat dry milk in PBS for 1 h at room temperature and incubated overnight at 4°C with primary antibodies. After washing, membranes were incubated with horseradish peroxidase (HRP)-conjugated secondary antibodies for 1 h at room temperature. Immunoreactive bands were visualized using enhanced chemiluminescence (ECL) and imaged with a ChemiDoc XRS+ Imaging System (Bio-Rad). Protein intensities were quantified in Image Lab (v6.0.1 build34, Bio-Rad). Data were normalized to GAPDH or another appropriate loading control on the same membrane. See Supplemental Table 5 for the detailed antibody information.

### Multi-electrode array recordings

Neuronal network activity was recorded using 48-well MEA plates containing 16 electrodes per well (Axion Biosystems). Before plating, wells were coated with 0.1 mg/mL poly-D-lysine (Sigma) and 20 ng/mL laminin (Thermo Fisher). Primary mouse glial cells (40,000 cells per well) were first seeded onto the recording area, followed 3 days later by 60,000 iGlut neurons. Glial cells and neurons were each plated as 10-15 μL droplets centered over the electrode array. After cell attachment, cultures were maintained as described above.

Spontaneous neuronal activity was recorded for 10 min daily beginning 7 days after neuronal plating. Recordings were acquired and spike metrics were extracted using AxIS software (Axion Integrated Studios, Axion Biosystems) with the following settings: 200-3,000 Hz band-pass filtering; spike detection threshold of 6 x SD; burst detection with a maximum interspike interval (ISI) of 100 ms and a minimum of 5 spikes; network burst detection with a maximum ISI of 100 ms, a minimum of 50 spikes, and a minimum percentage of participating electrodes (PE) of 35%; and synchrony analysis using a 20-ms window. To minimize the influence of plating variability, only wells with more than 12 active electrodes at day 32/34 were included in the final data analysis.

### Whole-cell current-clamp recordings in iGlut neurons

Whole-cell recordings from neuronal cell bodies were performed in current-clamp mode using a MultiClamp 700B amplifier (Molecular Devices). Patch pipettes were fabricated from borosilicate glass (World Precision Instruments) and had resistances of 3.0-4.0 MΩ when filled with an internal solution containing (in mM): 120 K-MeSO_4_, 10 KCl, 10 HEPES, 5 Na_2_-phosphocreatine, 4 Mg-ATP, and 0.4 Na-GTP, adjusted to pH 7.35 and osmolarity 280 mOsm/kg. The bath solution contained (in mM): 155 NaCl, 3.5 KCl, 1 MgCl_2_, 1.5 CaCl_2_, 10 HEPES, and 10 dextrose, adjusted to pH 7.35.

Resting membrane potential was measured immediately after break-in to the whole-cell configuration to minimize the influence of intracellular solution dialysis. Based on our recordings, substantial equilibration of the intracellular solution occurred within approximately 3-5 min. Action potentials for input-output curves were evoked by 500-ms current steps ranging from −30 to 210 pA, applied from either the native resting membrane potential or a holding potential of −65 mV. For action potential firing and waveform properties, only action potentials with a peak voltage > 0 mV and an upstroke velocity > 5 mV/ms were included. Action potential threshold was defined as the voltage corresponding to the peak of the second derivative of the action potential waveform. Action potential amplitude was measured as the peak voltage of the waveform, and half-width was measured at half-maximal amplitude. The fast afterhyperpolarization (fAHP) was defined as the maximum absolute hyperpolarization following the action potential. Upstroke and downstroke velocities were calculated as the maximum and minimum of the first derivative of the membrane potential, respectively. Input resistance was calculated from the voltage response to a 10-pA current step. Cells with input-resistance ≤ 100MΩ, or series resistance ≥ 20MΩ were excluded from analysis.

### Whole-cell voltage-clamp recordings in HEK cells

Whole-cell voltage-clamp recordings were performed on transiently transfected HEK293T cells 24-48 h after transfection. Transfected cells were identified by EGFP (expressed bicistronically via an internal ribosome entry site^6^) or mCherry fluorescence (expressed as a C-terminal fusion tag to *KCNH1*) using an Olympus IX73 inverted microscope equipped with a x40 objective. Patch pipettes were pulled from borosilicate glass capillaries (WPI) using a P-1000 micropipette puller (Sutter Instrument), with tip resistance of 3-4.5 MΩ when filled with the intracellular solution contained (in mM): 130 KCl, 10 HEPES, 7.5 Na_2_-phosphocreatine, 2 Mg-ATP, 5 EGTA, and 0.2 CaCl_2_, adjusted to pH 7.33 with ∼17.5 mM KOH and osmolarity adjusted with sucrose to 285-290 mOsm/kg. The extracellular solution contained (in mM): 140 NaCl, 3 KCl, 1.2 MgSO_4_, 2.4 CaCl_2_, 10 HEPES, and 20 glucose, adjusted to pH 7.37 with ∼4.5 mM NaOH and osmolarity adjusted to 300 mOsm/kg with sucrose.

Recordings were acquired at room temperature using an Axopatch 200B amplifier (Molecular Devices), filtered at 5 kHz, and sampled at 25 kHz. The amplifier feedback gain was set to 0.5 mV pA^−1^. Hardware leak subtraction was applied using at 1 GΩ resistive model, and series resistance was continuously monitored but not electronically compensated. Whole-cell currents were evoked from a holding potential of −90 mV using a 750 ms hyperpolarizing prepulse to −120 mV, followed by 1,000 ms test pulses from −80 to +50 mV in 10 mV increments and a 500 ms tail pulse to −30 mV, with a 12 s inter-sweep interval. Steady-state current density was calculated as the mean current during the final 10 ms of each test pulse normalized to whole-cell membrane capacitance. Voltage dependence of activation was assessed by calculating conductance at each test voltage as G(V) = I_ss_/(V-E_k_), where I_ss_ was the steady-state current measured over the final 10 ms of each test pulse and E_k_ was estimated as −99.4 mV using the Nernst equation. Conductance values were normalized to the maximum conductance within each cell, and the resulting conductance-voltage relationship was fitted with a Boltzmann function, G/G_max_ = 1/[1+exp (-(V-V_1/2_)/k)], to determine the half-maximal activation voltage (V_1/2_) for each cell.

### Knock-in mouse model

*Kcnh1^R357Q/+^*mice (hereafter *Kcnh1^Q/+^*) were generated by CRISPR/cas9 genome editing by the Northwestern University Transgenic and Targeted Mutagenesis Laboratory (TTML). CRISPR reagents were microinjected into C57BL/6J (B6) embryos at the two-cell stage. Potential founders were screened by Sanger sequencing of PCR products amplified using primers located outside the repair oligo homology region. Mosaic founders were bred to C57BL/6J inbred mice (Jackson Lab #000664) to generate N1 offspring, which were genotyped by Sanger sequencing to identify germline transmission of the R357Q allele. Potential off-target sites with fewer than three mismatches were PCR-amplified and analyzed by Sanger sequencing. Heterozygous N1. *Kcnh1^Q/+^* mice without detectable off-target edits were selected for subsequent breeding to establish the *Kcnh1^em1Kea^* line [MGI:8395115]. Because *Kcnh1^Q/+^* mice develop severe seizures and juvenile lethality, mice were maintained on chow compounded with the sodium channel blocker GS967 (8 mg GS967/kg chow; Research Diets, New Brunswick, NJ)^29^. To improve breeding performance and maternal tolerance of the neonatal ICV injection procedure, *Kcnh1^Q/+^* males were progressively backcrossed to 129S6/SvEvTac inbred females (129SVE, Taconic Biosciences). Experimental mice were on a mixed 129/B6 genetic background (generations ∼N3-N5), and male and female mice were used in all experiments.

Mice were genotyped by PCR followed by restriction fragment length polymorphism (RFLP) analysis. Genomic DNA isolated from tail biopsies was amplified using genotyping primers (forward, 5’-CAGTGCCTCTAATGCCAGTTC; reverse, 5’-CATTAAACTGGTATGGAGTGCC), followed by digestion with PvuII (#R0111, New England Biolabs) at 37°C for at least 1 h. PvuII digestion generated 492- and 50-bp fragments from WT and 492-, 313-, 179-, and 50-bp fragments from *Kcnh1^Q/+^*. Mice were maintained in a specific pathogen-free barrier facility on a 14-h light/10-h dark cycle with food and water available ad libitum. All animal procedures were approved by the Northwestern University Animal Care and Use Committee and conducted in accordance with the National Institutes of Health Guide for the Care and Use of Laboratory Animals.

### Intracerebroventricular administration of ASOs

Litters were randomly assigned to treatment groups. mASO-2 was diluted to the desired concentration in sterile D-PBS. Postnatal day 0-1 pups were cryo-anesthetized and freehand injections were performed using a 5 µL microsyringe (Hamilton, 65460-03) to deliver 4 µL of mASO-2 or vehicle (D-PBS) to the lateral ventricle. Following injection, pups were rewarmed and returned to the dam once they regained spontaneous movement.

### *Kcnh1* transcript Analysis

Total RNA was extracted from mouse brain tissue using TRIZol reagent (Invitrogen) according to the manufacturer’s instructions. First-strand cDNA was synthesized from total RNA using oligodT primers and Superscript IV (Life Technologies). Droplet digital PCR (ddPCR) was performed using ddPCR Supermix for Probes (No dUTP; Bio-Rad) with TagMan Gene Expression Assays for mouse *Kcnh1* (FAM-MGB, Mm01316769_m1) and the normalization controls *Tbp* (VIC-MGB, Mm00446971_m1). Reactions were partitioned using a QX200 Droplet Generator (Bio-Rad) and amplified using the following PCR conditions: 95 °C for 10 min; 44 cycles of 95 °C for 30 s and 60 °C for 1 min, with a ramp rate of 2 °C/s; followed by 98 °C for 5 min. Following amplification, droplets were analyzed using a QX200 Droplet Reader and QuantaSoft vl.6.6 software (Bio-Rad). *Kcnh1* transcript abundance was normalized to *Tbp*.

### Survival and behavioral assays

At postnatal day 22-24 (P22-P24), mice were weaned and put under continuous video monitoring with daily checks for morbidity or mortality. Mice with a body condition score of 2-1 were euthanized and counted as a death event^30^. Mice that were removed from the survival study for tissue collection or non-mutation related reasons were censored. Survival was compared between groups using the Mantel-Cox Log rank test with GraphPad Prism (v11.0.2). Video records were reviewed for instances of tonic-clonic seizures (bilateral convulsion with rearing and loss-of-posture) and tonic seizures (sudden, rigid stiffening with loss of posture and hindlimb extension at 180-degrees relative to the torso; may be followed by rhythmic jerking). For mice that died during video monitoring, 6 hours of video records prior to time of death were reviewed. For mice that survived, we reviewed 6 hours of video prior to the average time of death following weaning from GS967 (6.3 ± 0.7 days; n=30).

### Information for primers, antibodies, and ASOs

Primers were designed using Primer3^31^ and tested for gene specificity and primer efficiency before use. Primary and secondary antibodies were obtained from various vendors. The KCNH1 antibody for immunoblotting was verified by overexpression and siRNA-mediated knockdown experiments. Other antibodies were previously verified in our or other studies. ASOs were from IDT, with 2-MOE and PS modifications. ASOs for mouse injections had HPLC purification combined with Na+ salt exchange. All the sequences and other information were listed in the Supplemental Tables 1, 4, and 5.

### Data and statistical analysis

Data and statistical analyses for mouse studies were performed using GraphPad Prism (v11.0.2). All the other experiments were analyzed in R (v4.5.1). Details on statistical analyses, including sample sizes, tests performed, and multiple tests correct, if necessary, are provided in the figures or figure legends.

